# Optogenetic control of RelA reveals effect of transcription factor dynamics on downstream gene expression

**DOI:** 10.1101/2022.08.03.502739

**Authors:** Lindsey C. Osimiri, Alain R. Bonny, Seesha R. Takagishi, Stefanie Luecke, Nina Riehs, Alexander Hoffmann, Hana El-Samad

## Abstract

Many transcription factors (TFs) translocate to the nucleus with varied dynamic patterns in response to different inputs. A notable example of such behavior is RelA, a subunit of NF-κB, which translocates to the nucleus with either pulsed or sustained dynamics, depending on the stimulus. Our understanding of how these dynamics are interpreted by downstream genes has remained incomplete, partly because ubiquitously used environmental inputs activate other transcriptional regulators in addition to RelA. Here, we use an optogenetic tool, CLASP (controllable light-activated shuttling and plasma membrane sequestration), to control RelA spatiotemporal dynamics in mouse fibroblasts and quantify their effect on downstream genes using RNA-seq. Using RelA-CLASP, we show for the first time that nuclear translocation of RelA, without post-translational modifications or activation of other transcriptional regulators, is sufficient to activate downstream genes. Furthermore, we find that TNFα, a common endogenous input, regulates many genes independently of RelA, and that this gene regulation is different from that induced by RelA-CLASP. Genes responsive to RelA-CLASP show a wide range of dynamics in response to a constant RelA input. We use a simple promoter model to recapitulate these diverse dynamic responses, as well as data collected in response to a pulsed RelA-CLASP input, and extract features of many RelA-responsive promoters. We also pinpoint many genes for which more complex models, involving feedback or multi-step promoters, may be needed to explain their response to constant and pulsed TF inputs. This study introduces a new robust tool for studying mammalian transcriptional regulation and demonstrates the power of optogenetic tools in dissecting the quantitative features of important cellular pathways.

## Introduction

Transcription factors (TFs) are critical intracellular messengers that transmit to the genome information about the internal and external conditions of the cell. Many studies in the last decades have delineated how changes in nuclear TF concentration and post-translational modifications modulate the effect of TFs on their cognate genes (*1*–*4*). Increasing evidence shows that many TFs, such as NFAT and RelA, can additionally regulate their spatiotemporal dynamics in response to environmental inputs, and hence might encode information in these dynamics (*5, 6*).

Previous studies have used a variety of environmental and optogenetic inputs to regulate temporal dynamics of transcriptional regulators (*7*–*15*). A subset of these studies has also measured the effects of these dynamics on downstream gene expression, using techniques such as RNA-seq and reporter genes. For example, a study used gamma irradiation to induce pulses of p53, and then measured expression changes across the genome over a period of 12 hours using RNA-seq. By combining these data with ChIP-seq data, the authors showed that using a single dynamic input, p53 can activate downstream genes with different temporal patterns due to differences in TF-promoter binding kinetics (*10*). Another study regulated NFAT dynamics through optogenetic control of calcium concentration. With this method, it was found that a synthetic reporter did not differentiate between NFAT nuclear translocation dynamics, but instead activated in proportion to the integral of the NFAT nuclear localization (*7*). Finally, a recent study used microfluidics to precisely control the concentration of Tumor Necrosis Factor-ɑ (TNFɑ) delivered to cells, which then regulated the amplitude of NF-κB pulses of nuclear localization. As a result, the authors demonstrated that downstream genes had differential responses to pulsed inputs of different amplitudes (*8*).

A commonality across these studies is that they all modulate TF dynamics by controlling upstream regulators of the TF. As a consequence, such manipulations are pleiotropic, affecting many other downstream effectors in addition to the TF of interest. This can confound conclusions about the precise relationship between the dynamics of the TF and its target genes. As an example, a recent study quantified the effects of lipopolysaccharide (LPS) and TNFɑ using an *Ifnar*- and *Nfkbia*-knockout cell line. With these genetic modifications, these two inputs generated similar NF-κB translocation dynamics, thereby allowing authors to model stimulus-specific effects. Despite the similar NF-κB dynamics, hundreds of genes were differentially regulated by these stimuli, highlighting the complex interactions between different pathways in response to environmental inputs (*16*).

To dissect this complexity and elucidate the precise, quantitative effects of TF spatiotemporal dynamics on downstream gene expression in mammalian cells, we modify the yeast CLASP system for use across multiple cultured cell lines (*17*). With mammalian CLASP, we demonstrate optogenetic control of nuclear translocation for several TFs, including NFAT1 and RelA. We focus on RelA-CLASP and modulate its dynamics by inducing cells with pulsed and constant light inputs, thereby mimicking the effects of TNF and LPS, respectively. We monitor the effects of these TF spatiotemporal dynamics by measuring expression of downstream genes using RNA-seq. We observe that RelA-CLASP can activate downstream genes through its translocation alone, and that these genes have varied expression dynamics in response to a constant light input. We analyze these data using a simple model of promoter activation and gene expression. The dynamics of many genes could be captured with this model, which we use to extract quantitative relationships between model parameters. Additionally, this model can be used to predict the response of many genes to a pulsed light input, which we then experimentally validate with additional RNA-seq data. While many genes conform to this model, others show more intricate behaviors that highlight the complexity of signal processing that occurs downstream of RelA. Overall, our study introduces a highly effective optogenetic system in mammalian cells, and uses it to elucidate the RelA regulon and dissect promoter decoding of RelA translocation dynamics. These results add a deeper mechanistic understanding of the role of differing RelA translocation dynamics in response to environmental inputs.

## Results

### Designing Mammalian CLASP as a modular optogenetic tool

We have previously built, optimized, and demonstrated the utility of CLASP, a blue-light-responsive optogenetic tool, for regulating nuclear translocation of various transcription factors in yeast (*17*). Briefly, CLASP consists of two LOV2 domains. One LOV2 domain is tagged to the plasma membrane (pm-LOVTRAP), which sequesters a protein cargo tagged to the second LOV2 domain (Zdk1-cargo-yeLANS) in the dark (*18, 19*). Blue light stimulation activates CLASP by releasing the Zdk1-cargo-yeLANS from sequestration by pm-LOVTRAP at the plasma membrane and also by opening the yeLANS to reveal the nuclear localization sequence (NLS). The activation of both LOV2 domains leads to nuclear translocation of Zdk1-cargo-yeLANS. To extend these results to mammalian cells, we first engineered a CLASP DNA construct that would express readily across multiple cell lines, and would work across cell lines. To improve plasma membrane targeting for the LOVTRAP in mammalian cells, we swapped the Hs_RGS2 plasma membrane sequence used in yeast for a targeting sequence derived from Lyn kinase (*20, 21*). We next proceeded to optimize the method of delivery of CLASP DNA constructs. We created a three-plasmid system to deliver CLASP to mammalian cells: a plasmid expressing a fluorescent nuclear marker, a plasmid expressing the fluorescently-tagged pm-LOVTRAP system, and a plasmid expressing the fluorescently-tagged Zdk1-cargo-yeLANS protein (Figure S1A). These plasmids are delivered through lentiviral integration and can be transduced simultaneously or in series, depending on the goal of the user.

A critical benefit of the multi-plasmid system is that it allowed us to create a method that was more amenable to rapid screening of different cargo constructs across multiple cell lines. We created “chassis” cell lines (which we also refer to as pm-LOVTRAP cell lines) in HEK293T, 3T3, and MCF10A cells by transducing only the nuclear marker and pm-LOVTRAP plasmids; these could be used to screen any Zdk1-cargo-yeLANS construct of interest (Figure S1B). These chassis cell lines can be used with transient transfection to test whether a given cargo can be effectively sequestered and translocated by CLASP. With this three-plasmid system, a variety of cargos, including p53 and NFAT1 could be translocated to and from the nucleus in various cell lines with CLASP (Figures S1C-D). These experiments demonstrated that CLASP can be successfully transduced in every cell line that we have tried, and used to regulate nuclear translocation for a variety of transcription factor cargos. In summary, we have engineered CLASP to be a robust, modular tool for controlling nuclear translocation of a variety of protein cargos across mammalian cell lines.

### Quantification of RelA-CLASP response to light reveals reversible and dose-responsive dynamics

Given the modular nature of CLASP, we were particularly interested in testing its ability to control RelA (also known as p65), which is a transcriptional activator and subunit of the NF-κB complex. No previous reports have documented direct optogenetic control of RelA, despite its importance to cellular physiology. RelA regulates many cellular pathways, such as survival and inflammation, and is canonically studied for its response to immune system stimuli. In wildtype cells, the IκB proteins sequester RelA in the cytoplasm in the absence of stimulus. When cells are subjected to an environmental stimulus such as TNFɑ, IκB kinase (IKK) phosphorylates the IκB proteins, which leads to their degradation and the release of RelA (Figure 1A, top panel) (*6*). Critically, RelA is known to translocate to the nucleus with varied dynamics, undergoing repeated pulses or prolonged localization, depending on the stimulus (*8, 16, 22*). Optogenetic control of RelA translocation would therefore enable dissection of the effect of its nuclear translocation dynamics on downstream genes.

**Figure 1.**
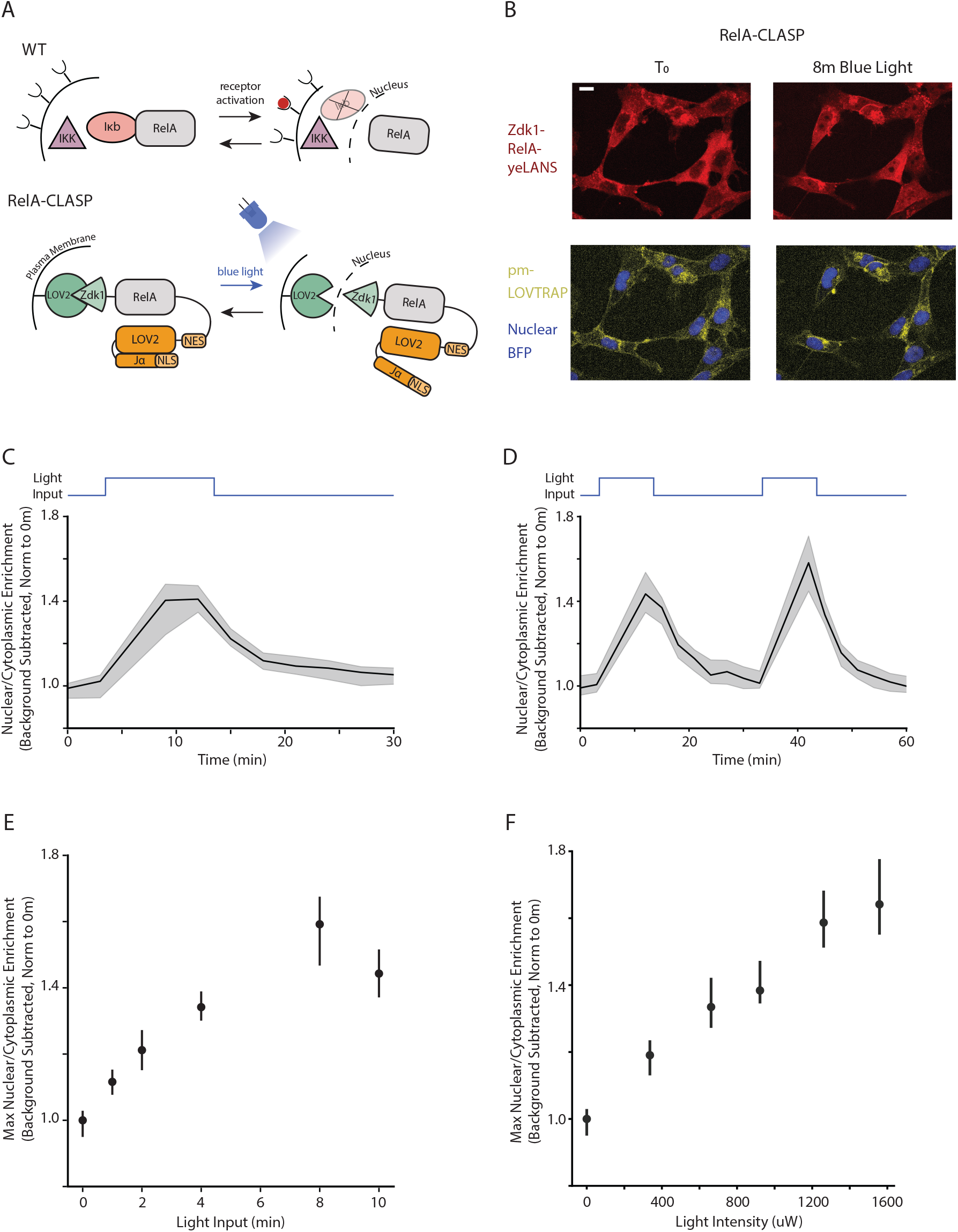

To maximize the chance of successful control over RelA nuclear localization, we expressed the RelA-CLASP construct in a 3T3 cell line that contains knockout mutations for RelA and three regulators of RelA translocation, the Inhibitor of nuclear kappa light chain gene enhancer in B-cells (IκB) proteins: IκBɑ, IκBβ, and IκBε (knockout cell line hereafter referred to as 3T3 *Nfkbia*^-/-^ *Nfkbib*^-/-^ *Nfkbie*^-/-^ *RelA*^-/-^). This cell line, which we termed RelA-CLASP cells, was therefore an ideal chassis for testing the ability of CLASP to regulate nuclear translocation of RelA. We also added several mutations to the sequence of *Mus musculus* RelA which were predicted to inactivate the nuclear localization sequence, thereby improving optogenetic control of RelA translocation (*23*–*25*).

Using this construct, we observed rapid blue light-mediated translocation of RelA from the plasma membrane to nucleus as measured by confocal microscopy; within 5 minutes of stimulation with blue light, nuclear translocation of RelA was evident (Figure 1B). Quantification of the response to blue light stimulation showed that nuclear translocation reached its maximum in less than 10 minutes and was reversible with nuclear exit occurring within 10-20 minutes after cessation of blue light (Figure 1C). Furthermore, consecutive pulses of blue light separated by 20 minutes generated pulses of RelA-CLASP nuclear translocation with similar dynamics and amplitudes (Figure 1D).

RelA-CLASP was also able to differentiate light inputs of different lengths and intensities. Varying blue light induction from 1-10 minutes, we found that the nuclear/cytoplasmic enrichment of RelA-CLASP increased with as little as 1 minute of blue light, reaching its maximum for a pulse duration of 8 minutes (Figures 1E, S2A). Nuclear/cytoplasmic enrichment of RelA-CLASP also increased approximately linearly as a function of blue light intensity (measured after 15 minutes of induction) (0-1.56 mW, induced with the Optoplate-96) (Figures 1F, S2B) (*26, 27*). Taken together, these data show that RelA-CLASP can quickly (≤ 8 minutes) and reversibly translocate to the nucleus, and that the magnitude of translocation can be tuned with light intensity.

### RelA-CLASP is insensitive to environmental inputs

RelA is endogenously regulated by a plethora of environmental inputs, including TNFɑ, Interleukin-1β (Il-1β), and LPS. Each of these inputs is recognized by a different cellular receptor: TNFɑ is recognized by the Tumor Necrosis Factor Receptor (TNFR), IL-1β activates the Interleukin-1 Receptor (Il1R), while LPS binds the Toll Like Receptor 4 (TLR4) (*28*) (Figure 2A, top panel). TNFɑ and Il-1β are cytokines produced during an immune response, and LPS is an endotoxin present on gram-negative bacteria that can induce an immune response. Since all three inputs converge on RelA, we wanted to determine whether these inputs retain their ability to activate RelA translocation and could therefore overpower RelA-CLASP sequestration. First, we sought to confirm prior reports documenting that wild type *Mus musculus* RelA translocates into the nucleus in response to its endogenous inputs in NIH3T3 cells. To do so, we built control cell lines (RelA-mScarlet cells) that heterologously expressed TagBFP as a nuclear marker and a wildtype *Mus musculus* RelA fused to mScarlet. These modifications were made in an otherwise wildtype NIH3T3 cell line that had intact endogenous control over nuclear translocation of RelA.

**Figure 2.**
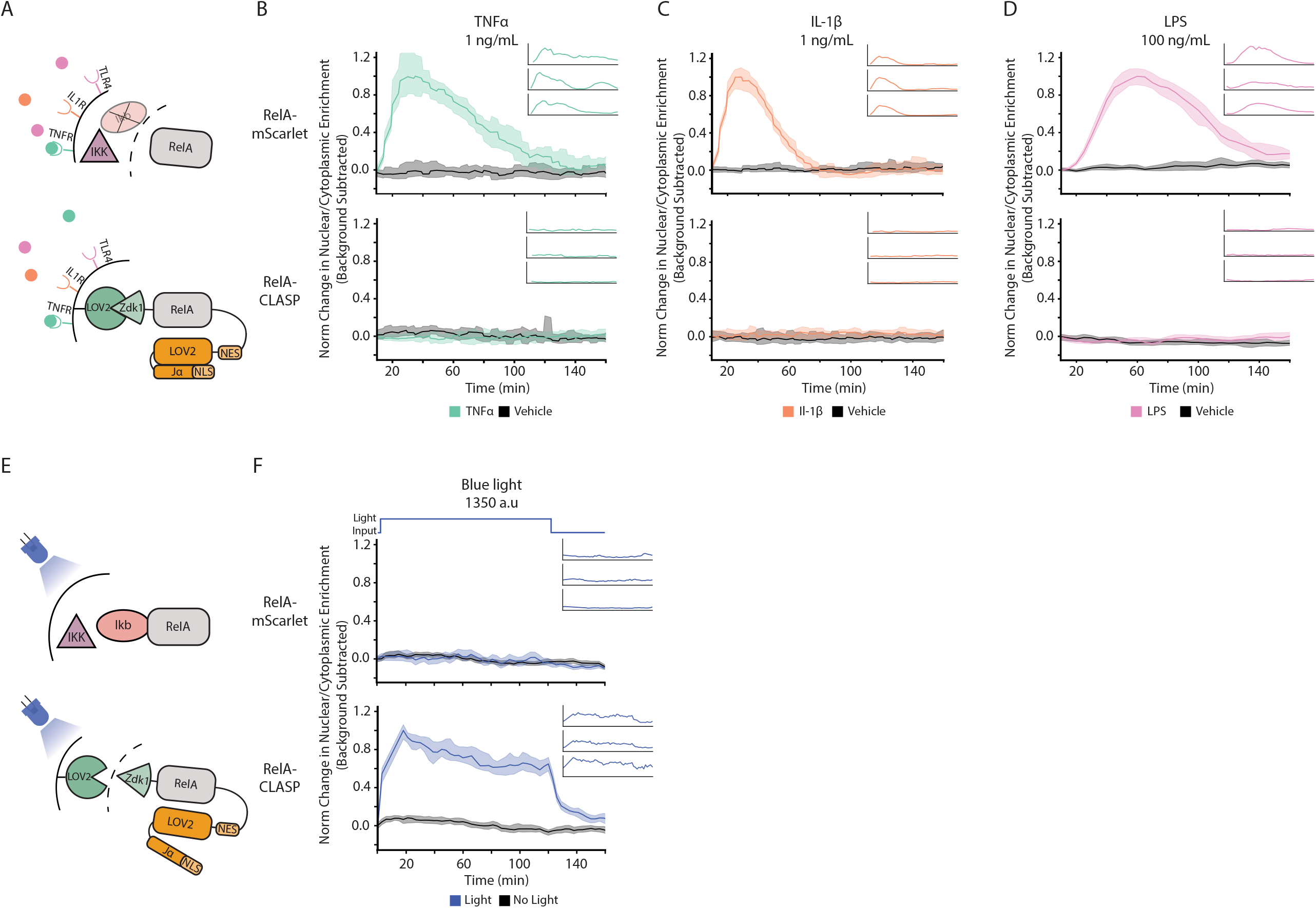

Prior work has shown that TNFɑ binding of the TNFR leads to activation of IKK, as well as other kinases like c-Jun N-terminal Kinase (JNK) (*28, 29*). Given the intact IKK-IκB pathway in RelA-mScarlet cells, RelA rapidly translocated to the nucleus upon stimulation with TNFɑ in this cell line (Figure 2B, green trace, top panel). This stimulation was specific to TNFɑ; addition of PBS + .1% BSA vehicle to RelA-mScarlet cells yielded no such response (Figure 2B, black trace, top panel). Previous studies have shown that TNFɑ induction leads to a coordinated first wave of translocation into the nucleus, followed by additional damped pulses (*8, 30*–*32*). Individual cell traces showed a similar finding, with some cells undergoing a short, single pulse within the first hour after induction, while other cells displayed two or more pulses of RelA translocation after induction (Figure 2B, inset, top panel). By contrast, in RelA-CLASP cells, nuclear/cytoplasmic enrichment in response to TNFɑ was minimal, and similar to that seen with vehicle induction (Figure 2B, green and black traces, bottom panel). Individual cell traces confirm that within the population, RelA-CLASP nuclear translocation was not induced by 1 ng/mL TNFɑ (Figure 2B, inset, bottom panel).

Il-1β activates the Il1R, which then signals through MyD88. This leads to activation of IKK, as well as other regulators like IL-1 receptor-associated kinase 1 (IRAK1) and JNK (*28*). Accordingly, RelA-mScarlet cells induced with 1 ng/mL of Il-1β responded with a coordinated translocation to the nucleus which resolved more rapidly than that induced by TNFɑ (Figure 2C, orange trace, top panel) (*16, 22*). Here again, there was minimal change in nuclear/cytoplasmic enrichment for RelA-CLASP cells induced with 1 ng/mL Il-1β (Figure 2C, orange traces, bottom panel).

LPS binds the TLR4, which activates the MyD88 pathway as well as interferon production (*28, 29*). Unlike the response to TNFɑ and Il-1β, RelA translocation in RelA-mScarlet cells in response to LPS was delayed, but extended for nearly two hours (Figure 2D, pink trace, top panel). Individual cell traces demonstrated that this prolonged translocation is uniform across cells, a key difference between the response to LPS and TNFɑ (Figure 2D, inset, top panel). Here again, despite the coordinated and prolonged response seen in RelA-mScarlet cells, translocation of RelA-CLASP in response to LPS was not distinct from translocation of RelA-CLASP after addition of vehicle (Figure 2D, bottom panel).

These data therefore indicate that RelA-CLASP is robustly sequestered and not induced by environmental stimuli. We therefore sought to probe next whether the complement is true -- that RelA itself is not responsive to light. To do so, we subjected both RelA-CLASP cells and RelA-mScarlet cells to a two-hour blue light input and measured nuclear localization (Figure 2E). When given a two-hour light input, RelA-CLASP maintained nuclear localization throughout the duration of the input, and then exited the nucleus shortly after the light input was turned off (Figure 2F, blue trace, bottom panel). Individual cell traces showed that within the population of RelA-CLASP cells, nuclear translocation and nuclear exit were synchronous across cells and also closely timed with the blue light input. For RelA-mScarlet cells, blue light did not lead to any appreciable nuclear translocation over the no light control (Figure 2F, blue and black traces, top panel).

Together, these data indicate that wild type RelA responds to environmental inputs by translocating to the nucleus, and that translocation of RelA-CLASP is exclusively controlled by blue light. While nuclear translocation of RelA is known to be necessary for expression of target genes, it is not known whether RelA translocation alone is sufficient for gene activation. As such, RelA-CLASP provided us with a unique opportunity to tackle this important question.

### TNFɑ and RelA-CLASP control gene regulons that overlap, but also differ in important ways

Many previous studies have used environmental inputs to regulate the nuclear translocation of RelA and quantify the effect of differential translocation dynamics on downstream genes (*8, 9, 13, 30*). As discussed previously, these inputs can have pleiotropic effects, activating many other pathways and regulators in addition to modulating RelA translocation. To quantify these pleiotropic effects for one such input, we delivered 1 ng/mL TNFɑ to pm-LOVTRAP cells, which express a nuclear marker and pm-LOVTRAP but not RelA. We then measured gene expression using RNA-seq at 0 hr, 1 hr, and 2 hr of TNFɑ induction (Figure 3A, top panel). In agreement with the idea that environmental inputs such as TNFɑ can elicit many cellular programs, we found that the expression of 1209 genes was significantly changed (FDR *p* < .05 as compared to 0 hr) by 1 hr or 2 hr TNFɑ induction, even in cells that lack RelA (examples shown in Figure 3B). Gene Ontology (GO) analysis of the 1209 genes modulated by TNFɑ indicated that pathways related to cell cycle, apoptosis, differentiation, and proliferation were significantly affected. However, terms related to NF-κB signaling, immunity, and inflammation were not significant in this analysis (Figure S3A).

**Figure 3.**
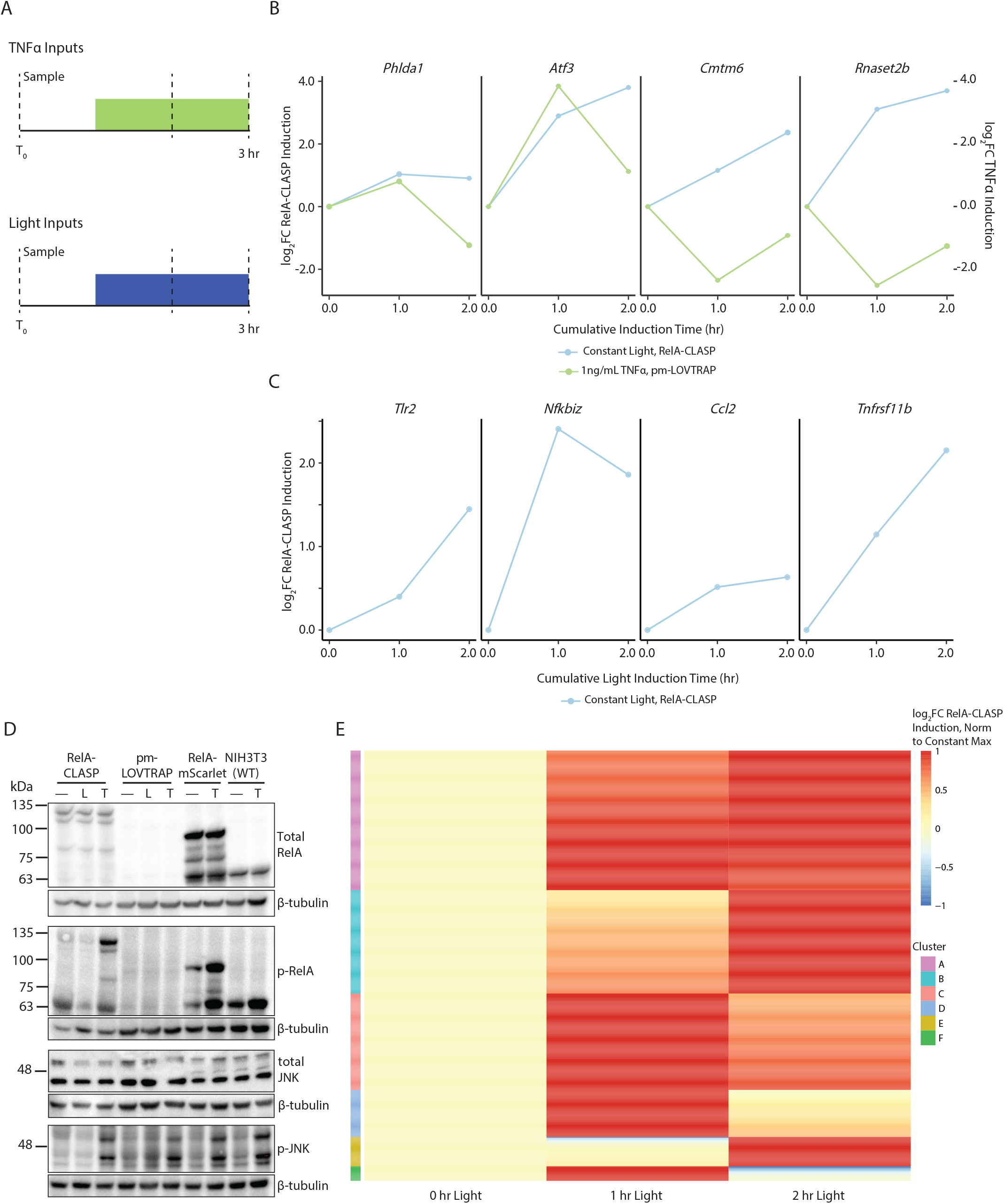

To obtain a global view of the ability of RelA nuclear translocation to induce gene expression, we next performed RNA-seq on RelA-CLASP cells induced with constant blue light at 1.38 mW (Figure 3A, bottom panel). We collected samples for sequencing at 0 hr, 1 hr, and 2 hr of blue light induction. As a control, we induced pm-LOVTRAP cells with the same light intensity and duration, and collected the same timepoints for sequencing. We sought to eliminate the effect of blue light by subtracting the log_2_ fold change in gene expression seen in pm-LOVTRAP from that seen in RelA-CLASP cells at each timepoint. We termed this metric log_2_FC RelA-CLASP induction (fold change is abbreviated as FC). As an additional control, we measured cell viability after exposure to 2 hours constant light at varied intensities. We found that pm-LOVTRAP and RelA-CLASP viability was not affected by the intensity of light used in our experiments (Figure S3C).

Overall, 650 genes out of the 1209 genes responsive to TNFɑ induction in pm-LOVTRAP cells were also significantly changed by constant light induction in RelA-CLASP cells (FDR *p* < .05 as compared to no input for the same cell line). This indicates that while these genes can be targets of RelA, they are also responsive to other signaling pathways. In general, these genes showed a more prolonged gene expression change in response to constant blue light in RelA-CLASP cells than in response to TNFɑ in pm-LOVTRAP cells. In fact, 585 out of these 650 shared genes are significantly induced at the 1 hr, but not at the 2 hr TNFɑ timepoint in pm-LOVTRAP cells. By contrast, only 124 of the 650 shared genes are not significantly induced at the 2 hr constant light timepoint in RelA-CLASP cells. Examples of these shared significantly regulated genes are plotted in Figure 3B. Genes such as *Phlda1* and *Atf3* reach similar maximal induction when induced by TNFɑ in the absence of RelA or constant RelA-CLASP input, but their induction proceeds with different dynamics. By contrast, others such as *Cmtm6* and *Rnaset2b* are induced by constant RelA-CLASP input but repressed by TNFɑ in the absence of RelA.

For the remaining 559 genes that are significantly changed by TNFɑ induction in pm-LOVTRAP cells but not by light in RelA-CLASP cells, GO analysis showed that they related to similar pathways as the entire set of genes significantly changed by TNFɑ induction (n=1209). Significant GO terms (*p* < .05) included response to tumor necrosis factor, ERK1 and ERK2 cascade, and epithelial cell proliferation (Figure S3B). These data indicate that TNFɑ, in the absence of RelA, induces a regulon that does not completely overlap with genes known to be downstream of RelA. This illustrates the wide-ranging changes caused by TNFɑ induction and points to the potential confounding effects of using environmental inputs to study the effect of nuclear translocation dynamics of a single transcriptional regulator.

### RelA-CLASP induced with a constant light input activates known RelA-responsive genes

To more specifically quantify the effects of RelA nuclear translocation dynamics on downstream genes, we analyzed the genome-wide effects of RelA-CLASP when induced with a two-hour constant light input as measured through RNA-seq. Differential gene expression analysis showed that 5034 genes were significantly regulated by RelA-CLASP in response to constant light (FDR *p* < .05 at 1 hr or 2 hr light input, when compared to 0 hr). Further examination of these genes showed that several were known to be targets of RelA (examples shown in Figure 3C). These genes included *Tlr2*, a Toll-like receptor that senses Pathogen-associated molecular patterns (PAMPs) and is activated in an NF-κB-dependent manner (*33*). *Nfkbiz* produces IκBζ, another inhibitor of RelA nuclear translocation; activation of IκB proteins, including IκBζ, acts as a feedback loop on RelA translocation (*34*). *Ccl2* is a chemokine induced in response to many inflammatory inputs, and *Tnfrsf11b* is a secreted protein that binds TNF-related apoptosis inducing ligand (TRAIL) and Receptor Activator of NF-κB Ligand (RANKL) (*35, 36*). The four genes regulated by RelA-CLASP translocation shown in Figure 3C had a log_2_FC RelA-CLASP induction varying from .5-2.5 over the course of 2 hours of constant light induction. Further, each of these genes displayed different dynamics of RNA expression despite the constant RelA-CLASP input.

Previous studies have shown that post-translational modifications of RelA, such as phosphorylation, are critical for downstream gene activation (*37*). Additionally, activation of RelA co-regulators, like JNK, CBP:p300, and IκBβ, in response to endogenous inputs has been shown to boost the ability of RelA to activate genes (*24, 38, 39*). Given that blue-light-induced RelA-CLASP translocation was capable of inducing many genes, we wondered if other RelA co-regulators were also induced by light. We turned to Western blotting to investigate this question. First, we confirmed the absence of endogenous RelA expression (65 kDa) in both RelA-CLASP and pm-LOVTRAP cells by blotting for total RelA (Figures 3D [top], S4A [top left], S4B [left], adjusted *p* < .01). We also confirmed that Zdk1-RelA-mScarlet-yeLANS (115 kDa) is expressed only in the RelA-CLASP cell line (Figures S4A [top left], S4B [right], adjusted *p* < .05). Second, we blotted for phosphorylated RelA and its co-regulator JNK, which are induced by environmental stimuli commonly used to induce RelA translocation (*8, 15, 16*). We observed no increase in phosphorylation of RelA and JNK in RelA-CLASP or pm-LOVTRAP cells after 30 min of blue light at 1.38 mW. As expected, 30 min of 1 ng/mL TNFɑ led to increased phosphorylation of RelA and JNK in RelA-CLASP, RelA-mScarlet, and wildtype NIH3T3 cells (Figures 3D [middle, bottom], S4A [bottom], S4C-D). Phosphorylated RelA and JNK were significantly lower in RelA-CLASP cells treated with blue light than those treated with TNFɑ (S4C-D, adjusted *p* < .001, adjusted *p* < .05, respectively). These data are the first demonstration that RelA is capable of controlling gene expression alone, without activation of its regulators. Indeed, these data provide direct evidence that nuclear translocation of RelA alone, in the absence of RelA phosphorylation, can induce known downstream genes.

### Genes regulated by RelA-CLASP show diverse dynamical behaviors

To further probe the quantitative dynamics of the genes responsive to RelA-CLASP under constant light stimulation (FDR *p* < .05, 5034 genes), we focused on those genes that had the top 15% of log_2_FC RelA-CLASP induction at either the 1 hr or 2 hr time points, thereby reducing the set of genes of interest to 896 genes. For comparison of the different dynamics of these genes without the confounding effect of their different induction amplitudes, we normalized the log_2_FC RelA-CLASP induction to its maximum for each gene. We then used longitudinal k-means clustering to group them based on quantitative differences in induction dynamics (*40*). Six clusters of dynamic gene expression trajectories emerged (Figure 3E, S3C). Cluster A (291 genes) reached peak induction at 1 hr of constant light, and maintained or slightly decreased induction at 2 hr of constant light. Cluster B (215 genes) increased in a graded fashion with each hour and peaked at 2 hr of constant light induction. Similar to cluster A, clusters C (201 genes) and D (98 genes) peaked at 1h of constant light, yet these genes had decreased induction by at least 15% (and up to 100%) of their maximum by 2 hr of constant light. Cluster E, on the other hand, represented 61 genes that induced minimally at 1 hr of constant light, and reached their peak at 2 hr of constant light. Cluster F (30 genes), similarly to clusters C and D, peaked at 1 hr of constant light induction, after which its log_2_FC RelA-CLASP induction decreased strongly to below 0.

These six clusters qualitatively represent four types of dynamic behaviors: early genes, proportional genes, late genes, and non-monotonic genes, a gene activation structure seen in previous studies (*10, 41, 42*). Early genes, like those in cluster A, are those that peak in log_2_FC RelA-CLASP induction at 1 hr of constant light and stay activated through 2h of constant light. Proportional genes, like those in cluster B, are those that increase proportionally in log_2_FC RelA-CLASP induction at both 1 hr and 2 hr. Late genes are not induced or may even be repressed at 1 hr of constant light induction, and are proceedingly induced at 2 hr of constant light induction; genes in cluster E fall into this category. Finally, non-monotonic genes are those that reach peak RelA-CLASP induction after 1 hr of constant light, and then decrease induction by 2 hr of constant light, which is seen in clusters C, D, and F. These data show a remarkable breadth of gene expression dynamics that can be induced by a constant two-hour input of RelA nuclear translocation.

### Genes that induce rapidly given a constant light input are predicted by a simple promoter model to generate varied responses to a pulsed light input

To understand the quantitative parameters that determine gene expression dynamics in response to a constant two-hour RelA-CLASP input, we built a simple model of mRNA expression. In this model, a promoter that exists in the OFF state (p_off_) transitions to the ON state (p_on_) with a rate that depends on RelA-CLASP nuclear concentration and the parameter k_on_. The promoter also transitions back to its OFF state with a constant off rate, k_off_. The state p_on_ is a transcription competent state that leads to mRNA production with a rate of β_1_. Transcription also proceeds at a basal rate β_0_. Finally, the mRNA decays with first order kinetics at a rate ɣ_1_ (Figure 4A). This simple model has been used in multiple investigations and shown to be able to reproduce a subset of gene expression dynamics (*16, 17*). However, it is incapable of recapitulating others, such as genes that respond slowly to constant TF input (late genes, cluster E in Figure 3E), as well as genes that peak early and then decline afterwards (non-monotonic genes, Clusters C, D, F in Figure 3E) because no such structural element (e.g. feedback loop) is present in the model. However, this simple model is sufficient to explain the dynamics of early (cluster A) and proportional (cluster B) genes, and therefore provides an opportunity to explore their quantitative parameters.

**Figure 4.**
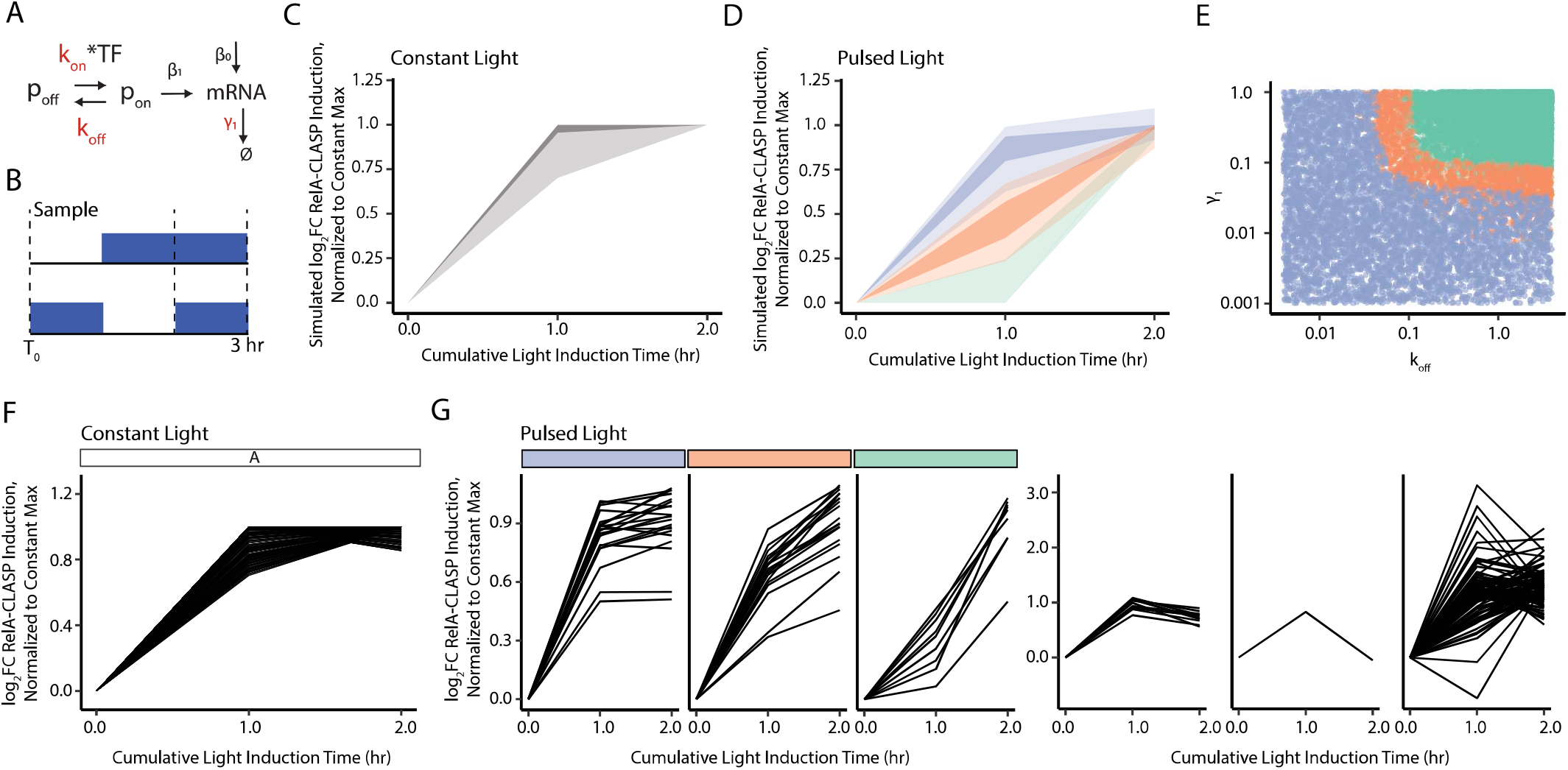

To do so, we simulated the model for 16,000 parameter sets varying k_on_, k_off_, and ɣ_1_. β_0_ and β_1_ were fixed across all parameter sets. We simulated mRNA time trajectories in response to a two-hour constant TF input for each combination of parameters. To allow comparison with the experimental data, we normalized these trajectories to the maximum simulated mRNA value, scored at either 1 hr or 2 hr after induction.

First, we filtered the simulated trajectories to only keep those that were within the data bounds seen in cluster A in the RNA-seq data (Figure 4C, F). Parameters were also filtered to remove those which led to lower log_2_FC RelA-CLASP induction than that seen in cluster A; these parameters were primarily those with low k_on_ and high k_off_. Examination of the parameter sets that could reproduce the early gene response showed that these parameters varied widely, but had a tradeoff between k_on_ and ɣ_1_ at the lower bound. That is, a lower k_on_ necessitated a higher ɣ_1_, and vice-versa (Figure S5A-C). ɣ_1_ is inversely proportional to basal mRNA expression. Accordingly, a higher ɣ_1_ leads to lower basal mRNA expression and faster dynamics, allowing parameter sets with any value of k_on_ to increase their log_2_FC RelA-CLASP induction at a speed commensurate with the data once the light turned on. On the other hand, parameters with a lower ɣ_1_ required higher k_on_, which increases the rate of mRNA expression in response to RelA input (Figure S5D).

Different combinations of model parameters (k_on_, k_off_, ɣ_1_) were responsible for these similar responses to a constant two-hour RelA-CLASP input. This behavior could be the result of parameter degeneracy, or it could also reflect genuine differences in the characteristics of the promoters. We hypothesized that we could demonstrate whether different parameters underlie the same response to prolonged RelA-CLASP translocation by subjecting these cells to a more dynamic light input. If the genes exhibit a homogenous response to the dynamic input, then it is likely that parameter degeneracy exists in the model. However, if the input yields different gene expression responses that relate to subsets of parameter combinations, then it is possible that these promoters possess different parameters that can only be revealed with more dynamic inputs.

We first used the computational model to test this hypothesis, simulating the response to a pulsed RelA-CLASP input for the same parameter sets previously identified as yielding an early gene response to a constant input. This input delivered light for a cumulative 2 hours, but with two hour-long pulses separated by one hour of no light (Figure 4B). Despite the uniform response to constant light inputs, the simulated responses to this pulsed input exhibited a variety of behaviors. In order to segment these dynamic responses, each simulated gene expression trajectory was normalized to its maximum value for the pulsed input to remove the effect of magnitude and then clustered using k-means longitudinal clustering (*40*). We identified three clusters as the most parsimonious division. To facilitate comparison with the plots showing response to constant light input, we normalized the simulated gene expression to the maximum simulated expression in response to a constant input (Figure 4D, S5E). The first cluster (purple) corresponded to parameter sets with a response that reached near-maximal induction at the first timepoint after light induction, that is, after one hour of TF input and a one hour OFF period (represented in Figure 4D as ‘1 hour cumulative light induction time’). These parameter sets are capable of inducing slightly more mRNA in response to a pulsed TF input than a constant input (Figure 4A). The second cluster (orange) contained parameter sets that responded commensurately to the pulsed TF input in log space, with expression increasing at each timepoint. Finally, the third cluster (green) consisted of parameter sets that had low gene expression at the first timepoint after light induction and reached maximal expression at the end of the experiment.

Examination of the parameter values that generated these different classes of dynamic trajectories in response to a pulsed input showed a clear pattern: the clusters were most differentiated by the relationship between k_off_ and ɣ_1_. The orange and green clusters (Figure 4D) were characterized by simultaneously higher k_off_ and ɣ_1_ values relative to all parameter sets that recapitulated the early gene response to a constant input (Figures 4E, S5F-G). On the other hand, the purple cluster (Figure 4D) contained parameter sets with low k_off_ and ɣ_1_, high k_off_ and low ɣ_1_, or low k_off_ and high ɣ_1_. Indeed, for both the orange and purple clusters, there was a tradeoff between ɣ_1_ and k_off_, where higher k_off_ required lower ɣ_1_ and vice-versa (Figure 4E). However, the purple cluster required either lower values of k_off_ or ɣ_1_ than those within the orange cluster, and could have parameter sets with simultaneously low k_off_ and ɣ_1_.

The importance of the relationship of k_off_ and ɣ_1_ can be explained through examination of the model. It was previously shown that the rate at which mRNA expression increases in response to a TF input is dependent on k_on_, k_off_, and ɣ_1_; therefore all three parameters affect mRNA expression while the light is ON (*17*). However, for the pulsed input, after the light turns OFF and RelA-CLASP exits the nucleus (within 15 minutes of the end of the light input), mRNA expression is largely determined only by k_off_ and ɣ_1_.

For the green cluster, high k_off_ and ɣ_1_ allowed log_2_FC RelA-CLASP induction to reach steady-state value during the first hour of the experiment, when the light is ON. During the OFF period, though, these parameters also yielded fast mRNA degradation and rapid promoter shutoff (Figure S5H). As a result, the log_2_FC RelA-CLASP induction measured at the first timepoint after light induction was low (Figure 4D). Still, due to the faster kinetics of the green cluster, log_2_FC RelA-CLASP induction reached its maximum value again at the end of the experiment.

By contrast, the purple cluster contained parameter sets with low k_off_ and ɣ_1_, high k_off_ and low ɣ_1_, or low k_off_ and high ɣ_1_. This caused, on average, a slightly slower rate of increase for log_2_FC RelA-CLASP induction during the first hour, when the light is ON. However, these parameters also had a slower rate of shutoff during the time that the light was OFF. For those parameters with low k_off_, the promoter continued to produce mRNA even in the absence of TF input. For those with low ɣ_1_, there was slow degradation of the mRNA. Both of these scenarios lead to sustained mRNA expression during the light OFF period, and high log_2_FC RelA-CLASP induction at the final timepoint (Figures 4D, S5H). Due to the mRNA accumulated before the second pulse, the purple cluster is able to generate slightly higher mRNA expression in response to a pulsed input at the final timepoint than a constant two-hour input.

Finally, the orange cluster showed an intermediate phenotype where, on average, log_2_FC RelA-CLASP induction increased more slowly than the green cluster, but more quickly than the purple cluster while the light was ON. However, the increased values of k_off_ and ɣ_1_ relative to the purple cluster also led to faster mRNA decay and promoter shutoff during the light OFF period (Figure S5H). This increased mRNA decay and promoter shutoff led to a moderate value of log_2_FC RelA-CLASP induction at the first timepoint after light induction, and maximal or near-maximal log_2_FC RelA-CLASP induction at the final timepoint.

We vetted the predictions generated through simulation of a pulsed TF input with an RNA-seq experiment using a pulsed light input (Figure 4B). For these data, we calculated log_2_FC RelA-CLASP induction for the genes identified in cluster A (Figure 4F) that were significantly induced (FDR *p* < .05) by pulsed light inputs and normalized each gene to its maximum log_2_FC RelA-CLASP induction in response to constant light. We then clustered the gene expression dynamics for the pulsed light input. Doing so identified 7 clusters, three of which closely resembled those predicted by the simple model (16.8% of genes in clusters A, Figure 4G). This strongly implied that these genes could be well-modeled by the most parsimonious simple promoter representation. However, there were still 4 clusters of genes that were not predicted by the simple model. The first two clusters of genes both peaked early in response to pulsed inputs, and then decreased induction during the second hour of light input (fourth, fifth graphs, Figure 4G; 4.1% of early genes). This decrease in induction in response to TF input cannot be predicted by the simple model. The third cluster consists of genes with a maximal induction that is much higher for pulsed light inputs than for constant light inputs, which can also not be predicted by any parameter set tested with the simple model (sixth graph, Figure 4G; 35.7% of early genes). The final cluster, which was not plotted, consists of genes not significantly induced (FDR p > .05) by pulsed light inputs (43.3% of early genes). This lack of induction in response to pulsed inputs additionally cannot be predicted by this model.

### Genes that induce gradually given a constant light input are predicted by a simple promoter model to yield varied responses to a pulsed light input

As discussed previously, the simple model can also be used to recapitulate the proportional response to constant TF inputs in log space. 16,000 parameter sets varying k_on_, k_off_, and ɣ_1_ were generated as described before, and then used to simulate mRNA dynamics in response to a constant RelA-CLASP input (Figure 5A). Each of these simulations was normalized to the maximum mRNA value simulated, and then all parameter sets were filtered for those that generated gene expression dynamics within the bounds of the proportional genes (cluster B, Figure 3E) identified by the RNA-seq data (Figure 5B). Overall, these parameter sets had lower ɣ_1_ and lower k_on_ values than the full set of parameters tested (Figure S6A-C). Lower k_on_ values caused mRNA expression to rise more slowly (*17*). Furthermore, lower ɣ_1_ meant that mRNA decay occurs slowly, allowing the mRNA to accumulate over the length of the constant TF input (Figure S6D).

**Figure 5.**
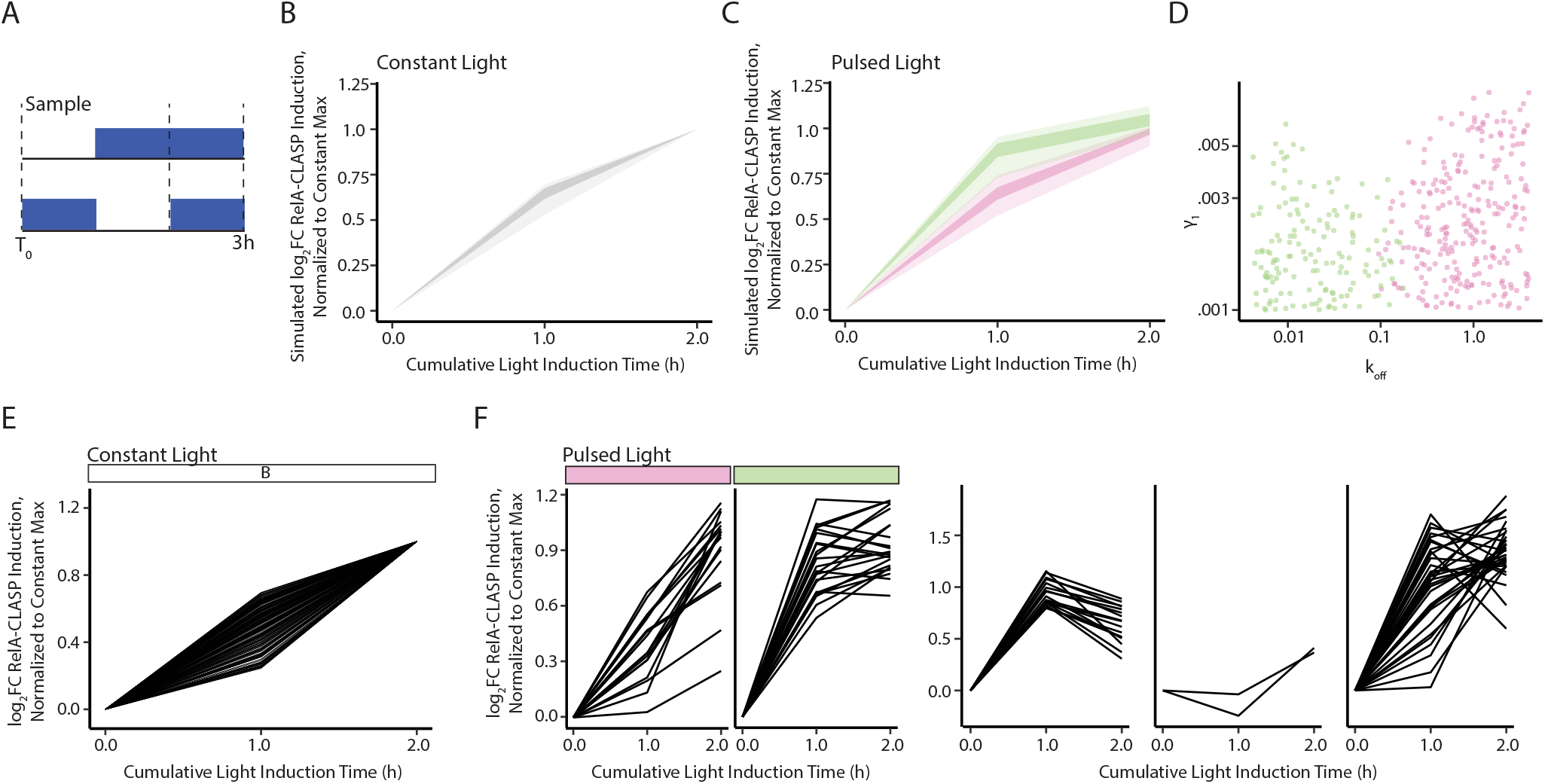

Next, we used these parameter sets to simulate the model response to a pulsed RelA-CLASP input that delivered a cumulative 2 hours of light, but in one hour pulses separated by an hour of light OFF (Figure 5A). We again clustered the simulated responses after normalizing each trajectory to its maximum value in response to the pulsed TF input. To facilitate comparison across plots, we plotted both clusters with simulated log_2_FC RelA-CLASP induction normalized to its maximum value for the same parameter set when simulated with a constant two-hour input. Clustering identified two qualitatively different responses to a pulsed RelA-CLASP input (Figure 5C, S6E). The green cluster showed a steeper rise in log_2_FC RelA-CLASP induction at the first timepoint after light induction (given one hour ON and one hour OFF) than the pink cluster. These two behaviors also differed in their associated k_off_ values (Figure 5D, S6F). In particular, parameter sets in the pink cluster had higher k_off_ values relative to all parameter sets that generated a proportional increase in log_2_FC RelA-CLASP induction.

Many of the parameter sets in the pink cluster also had a slightly higher k_on_ (Figures 5D, S6F-G). As a result of the increased k_on_, these genes increased mRNA expression more quickly during light ON; however, the increased k_off_ led to rapid shutoff of the promoter during light OFF. For all promoters that generated a proportional response to constant TF inputs in log space, ɣ_1_ is necessarily low. Therefore, while log_2_FC RelA-CLASP induction slightly decreased during light OFF, it did not completely decay. During the second RelA-CLASP pulse, this cluster reached its maximal log_2_FC RelA-CLASP induction (Figure S6H). On the other hand, the parameter sets in the green cluster had lower k_off_ values (Figure 5D, Figure S6F-G). Lower k_off_ combined with the same low ɣ_1_ constraint resulted in mRNA levels that kept slightly increasing during the light OFF period. This resulted in a slightly higher log_2_FC RelA-CLASP induction at the first timepoint after light induction (Figure S6H).

Here again, we compared the predictions from these simulations with the RNA-seq data for pulsed inputs, by clustering the response to pulsed light inputs for the genes assigned to cluster B (Figure 5E). This resulted in 6 clusters, two of which qualitatively recapitulated responses predicted by the simple model (Figure 5F; 19.1% of proportional genes). The third cluster of gene trajectories displayed a strong induction at the first timepoint after light induction followed by decreased induction at the final timepoint (7.9% of proportional genes). The fourth cluster consisted of two genes that had a delayed response to pulsed light inputs. The fifth cluster displayed genes that activated much more strongly in response to pulsed light inputs than constant light inputs (14.9% of proportional genes). Lastly, many genes from cluster B (57.2% of proportional genes) were in fact not significantly (FDR *p* < .05) regulated by pulsed light inputs. These behaviors cannot be recapitulated by the model, indicating that for these genes, likely more sophisticated regulation is at play. Nonetheless, given that RelA canonically regulates genes in conjunction with other transcriptional regulators, and activates several feedback loops, we find it remarkable that a meaningful subset of its downstream genes can be modeled as simple promoters.

## Discussion

In this study, we extended our previous efforts to build a modular and reversible optogenetic tool to an additional model system, mammalian cells, and then used this tool to generate a novel transcriptomic dataset that measures the effect of RelA nuclear translocation on downstream genes. Importantly, we were able to provide the first direct demonstration that RelA translocation, without phosphorylation or activation of co-regulators, is able to induce downstream gene activation. Using a computational model of gene expression, we were able to further probe this dataset to categorize genes whose promoters might activate linearly in response to RelA nuclear concentration. We also identified promoters that cannot be modeled linearly, which suggests more complex features such as multistep activation or feedback.

To dissect the intricate solo gene regulon of RelA-CLASP, we asked whether the simple gene expression model could constrain parameter relationships of promoters that are explained by this model. For early genes, those that reach near-maximal log_2_FC RelA-CLASP induction within one hour of constant light, we found that the constant input could only mildly constrain the relationship between k_on_ and ɣ_1_. Notably, this two-hour continuous pulse of RelA was picked to mimic nuclear translocation in response to LPS. By contrast, a pulsed RelA input with two one hour pulses separated by one hour of light OFF was highly informative – it delineated three classes of promoters, differentiated by their k_off_ and ɣ_1_. This computational experiment was further corroborated with RNA-seq data that displayed the 3 quantitatively different classes of promoters, in addition to other classes.

For genes that proportionally increased their log_2_FC RelA-CLASP induction in response to constant light, the parameters were better constrained – they cannot have a large k_on_ or ɣ_1_. Furthermore, the gene expression response of these promoters for the pulsed input was more similar to the response following the continuous input, though with additional subtleties. This behavior was also similar to a subset of the early genes (cluster A) when subjected to the same pulsed input (compare for example pink in Figure 5C and orange in Figure 4D). Interestingly, this same behavior is generated by different underlying parameters. Therefore, using only the pulsed input to constrain the parameters of these quantitatively different promoters would have failed. These observations highlight the complex experimental design needed to unravel quantitative parameters. As a case in point, many of the genes that are responsive to RelA cannot be explained by a simple model and may require additional dynamic inputs to constrain their parameters. We hope this will be the subject of many future studies that move the field into quantitative and predictive understanding of promoter decoding of RelA translocation dynamics.

These investigations will require precise tools such as CLASP, rather than environmental inputs. Indeed, while the RelA-CLASP gene regulon overlaps with the TNFɑ-induced regulon, the two are not identical. For example, RNA-seq data in Figure 3 collected following TNFɑ induction in a RelA and IκB knockout cell line unambiguously show that TNFɑ regulates many genes independent of RelA. These data demonstrate the difficulty in using environmental inputs alone to study the downstream effects of TF dynamics. Future studies measuring the epistatic relationships between RelA and various environmental inputs, as well as the relationship with other transcription factors, would expand our understanding of RelA transcriptional activation.

The latter considerations are a clear limitation of this study. Combinatorial control of gene expression is widespread, if not ubiquitous, in mammalian cells. In future studies, CLASP could be composed with environmental inputs to precisely regulate RelA translocation during stimulus, thereby allowing other pathways to be activated at the same time. This would be possible because of the robust sequestration of pm-LOVTRAP and the knockout of IκB proteins, as seen in Figure 2. A recent study used chemical inputs and optogenetic control of Erk to discern the effect of Erk translocation on cell proliferation; future studies focusing on RelA could also elucidate the effects of dynamics in a similar fashion (*12*). Additionally, other studies have compared dynamics across stimuli to understand what features of a dynamic RelA input help the cell to differentiate between inputs (*43, 44*). RelA-CLASP could be used to further substantiate the computational analyses that were used to draw conclusions on how genes use parameters such as amplitude, pulse width, and frequency to differentiate inputs. Importantly, these relationships should also be delineated across additional cell lines, since cell identity can affect gene response (*13*).

This study is limited by the use of bulk RNA-seq data, which might obscure the individuality of response in single cells. This individuality could expand our quantitative understanding of promoter decoding of RelA transcription dynamics. Future experiments using RelA-CLASP could use single cell RNA-seq (scRNA-seq), single molecule RNA FISH (smFISH), or MS2 systems to further probe heterogeneity in response to the same dynamic TF input. Notably, several recent studies have used environmental inputs to modulate localization of transcriptional regulators like RelA and Erk, and then measured the heterogeneous response of single cells with these methods (*9, 32, 45, 46*). These studies observed that TF translocation dynamics, as well as downstream gene expression dynamics, are heterogeneous amongst single cells in response to environmental inputs. Tracing the roots of this heterogeneity, and ascribing it to the TF signal itself or signal processing at the level of the genes may be facilitated by a uniform dynamic TF input generated by optogenetic control.

## Materials and Methods

### Experimental details

#### Mammalian cell culture

HEK293T and LX293T cells were cultured in 1 g/L glucose DMEM (Life Technologies 11885076), 1% Antibiotic-Antimycotic (Thermo 15240062), and 10% Fetal Bovine Serum. NIH3T3 and 3T3 *Nfkbia*^-/-^ *Nfkbib*^-/-^ *Nfkbie*^-/-^ *RelA*^-/-^ cells were maintained in 1 g/L glucose DMEM (Life Technologies 11885076), 1% Antibiotic-Antimycotic (Thermo 15240062), and 10% heat-inactivated Bovine Calf Serum (UCSF Cell Culture Facility, HyClone, lot number AZM197696). Heat inactivation was accomplished by heating serum at 56°C for 30 minutes. After heating, serum was cooled to room temperature before media production. MCF10A cells were cultured in DMEM/F12 (Thermo 21331020), 5% Horse Serum (UCSF Cell Culture Facility), 0.1 mg/mL EGF, 4 mg/mL Insulin (Gibco 12585014), 1 mg/mL Hydrocortisone, .1 mg/mL Cholera toxin (Sigma C8052), and 1% Antibiotic-Antimycotic (Thermo 15240062). All cells were cultured at 37° C and 5% CO_2_. NIH3T3 and 3T3 *Nfkbia*^-/-^ *Nfkbib*^-/-^ *Nfkbie*^-/-^ *RelA*^-/-^ cells were passaged every three days; HEK293T, LX293T, and MCF10A cells were passaged every other day.

#### Plasmid and cell line construction

Hierarchical golden gate assembly was used to assemble all plasmids (*21, 47*). BsaI and BsmBI sites were removed from parts to enable further assembly. Parts were generated through PCR or ordered as gBlocks from IDT. Plasmids were grown and prepared from DH5ɑ, XL1 Blue, Mach1, or Stbl3 competent cells (Macrolab, Berkeley, CA). For lentiviral transduction, plasmids were first transfected into LX293T cells at 80% confluency using Lipofectamine 2000 (Thermo 11668019), the plasmid of interest, and two plasmids encoding second generation lentiviral envelope and packaging vectors (MDG.2 and CMV). Transfection reagent and media were removed from LX293T cells approximately 16 hours later and transfected cells were re-fed with 1 g/L glucose DMEM (Life Technologies 11885076), 1% Antibiotic-Antimycotic (Thermo 15240062), and 10% Fetal Bovine Serum. 24 hours later, the media was removed from transfected LX293T cells and filtered through a .45 micron filter to remove cell debris. For 3T3 *Nfkbia*^-/-^ *Nfkbib*^-/-^ *Nfkbie*^-/-^ *RelA*^-/-^ cells, polybrene was added to the filtered viral supernatant to achieve a final concentration of 4 µg/mL after adding to cells. Viral supernatant was then added to cells for transduction slowly on top of media. After addition of viral supernatant, cells were spun at 800xg for 45 minutes to increase transduction efficiency. After 16-24 hours of incubation with viral supernatant, cells were re-fed with fresh media. After transduction, cells were sorted to select the population of interest.

#### Cell selection via sorting

To prepare for sorting, cells were lifted using trypsin and resuspended in the corresponding media to quench trypsin activity. Afterwards, cells were spun down at 400xg for 5 minutes to form a pellet and placed on ice. This pellet was then resuspended in PBS for sorting. Sorting was performed on a BD FACSAria II. BFP was assessed using the BV405 channel (405 nm excitation, 450/50 nm filter), mScarlet was measured using the mCherry channel (561 nm excitation, 610/20 nm filter), and IRFP was assessed using the APC-Cy7 channel (633 nm excitation, 780/60 nm filter). Cells were sorted into fresh media and re-plated after sorting.

#### Modified RelA

Residues Lys^301^, Arg^302^, Lys^303^, Leu^438^, Leu^441^, and Phe^443^ in *M. musculus* RelA were mutated to alanines, and His^440^ was mutated to a glutamine.

#### RelA-CLASP cell line generation

RelA-CLASP was generated through lentiviral transduction of the bulk-sorted pm-LOVTRAP chassis cell line (background 3T3 *Nfkbia*^-/-^ *Nfkbib*^-/-^ *Nfkbie*^-/-^ *RelA*^-/-^) with the Zdk1-RelA-mScarlet-yeLANS construct (chassis cells were sorted for iRFP expression (pm-LOVTRAP) and nuclear marker expression (BFP)). After transduction, this cell line was further selected to generate a clonal cell line which is referred to as RelA-CLASP in this study. Single cells expressing BFP, low IRFP, and low RFP were sorted into a 96 well plate and then clonally expanded. After expansion, clonal cell lines were assessed for continued expression of fluorophores and responsiveness of CLASP construct. A single clonal cell line, termed RelA-CLASP F8 lo, was selected for use in this study.

#### Microscopy

For microscopy, a 96-well glass-bottom plate (Thermo Fisher 164588) or a 24-well microscopy plate (Ibidi 82406) was incubated with 0.1 mg/mL Poly-D-Lysine (Gibco A3890401) at room temperature for 1 hour, after which the plate was washed three times with sterile water and left to dry for 2 hours. After drying, cells were seeded at 8000 cells/well (96 well plate) or 35000 cells/well (24 well plate) in media without phenol red (Thermo Fisher 11054020) with glutamine supplemented (Life Technologies 35050-061). 48 hours later, cells were imaged. Microscopy for all figures (except for Figures 1B and S2D-E) was performed on an inverted Nikon Ti microscope equipped with a CSU-22 spinning disk confocal, EMCCD camera, and custom 4-line solid state laser launch. Imaging took place inside a cage incubator which maintained temperature, CO_2_, and humidity throughout the experiment. Images were taken using a 40x/0.95 objective, and cells were illuminated with 405, 561, and 640 nm lasers. For any images where cells are induced with light on this microscope, cells were covered with a BreatheEasy seal and a custom-printed Optoplate holder was mounted on top of the cells. The Optoplate was then placed on top of the holder to induce the cells. For timecourse microscopy, images were acquired every 3 minutes. For Figures 1B and S2D-E, microscopy was performed on an inverted Nikon Ti microscope with an Andor iXon Ultra DU888 1k x 1k EMCCD and Andor 4-line laser launch. An Oko stage was used to maintain temperature and atmosphere control. For these panels, cells were induced using 488 nm light produced by imaging GFP.

#### Drug induction

IL-1β (Peprotech, 211-11B) was diluted to a stock concentration of 100 ng/mL in sterile water with .1% Bovine Serum Albumin (BSA). LPS (Sigma, L2880) was diluted to 10 µg/mL in phosphate-buffered saline (PBS). TNFɑ (R&D Systems, 410-MT) was diluted to a 100 ng/mL stock solution in PBS with 0.1% BSA. All stocks were at 100X concentration. Prior to induction, stocks were diluted to 2X or 3X in media and then added to cells to a final 1X concentration (TNFɑ, IL-1β: 1 ng/mL, LPS: 100 ng/mL).

#### Light induction using Optoplate-96

Optoplate-96 was programmed using the OptoConfig-96 program (*48*). Cells were induced with up to 12 minutes of constant light input, followed by a pulsed light input of 2 seconds ON/2 seconds OFF to reduce blue light toxicity.

#### RNA-seq

Cells were seeded with 40,000 cells/well in 24 well plates (Ibidi 82406) 2 days prior to experiment, so that they would be 80% confluent when induced. Five replicate wells were seeded for each input and cell line. Immediately prior to experiment, cells were induced with vehicle (PBS + 0.1% BSA) or TNFɑ diluted in media to a 2X concentration. For induction, 0.5 mL of media was removed from the well and .5 ml of induction media was added. After induction, a BreatheEasy seal (Sigma Z380059) was placed on cell plate. For cells induced with TNFɑ for 1 hr, cells were placed into experiment incubator for 2 hours, removed, induced, and then removed 1 hour later for harvesting. For cells induced with TNFɑ for 2 hr, cells were placed in experiment incubator 1 hr before induction. For cells induced with light, cells were induced with vehicle media and placed into incubator with Optoplate-96 directly on top of BreatheEasy seal for 3 hours. All cells were harvested immediately after induction, and RNA was isolated from cell pellets using the Lexogen SPLIT RNA extraction kit. After extraction, RNA quality was assessed using the Agilent Pico RNA kit, and quantified using a Nanodrop. Following extraction, RNA samples were diluted using concentrations estimated by Nanodrop. Libraries were prepared using the Lexogen Quantseq 3’ RNA-seq Library Prep Kit on 250 ng RNA from each sample. Library quality was assessed using the Agilent High Sensitivity DNA Kit, and quantity was measured using the Qubit dsDNA HS Assay Kit. Libraries were diluted to equimolar concentrations and pooled. Pools were subject to single-end sequencing on the Illumina HiSeq 400.

#### Western blot

For Western Blotting analysis of protein levels, cells were seeded as described for RNA-seq. Immediately prior to experiment, cells were induced with vehicle (PBS + 0.1% BSA) or TNFɑ diluted in media to a 3X concentration as described for RNA-seq. Cells induced with vehicle were then given no light input or 1.38 mW blue light for 30 minutes. Cells induced with TNFɑ were induced with a final concentration of 1 ng/mL TNFɑ for 30 minutes. For whole cell lysis, media was removed, the cells washed once with cold PBS, and then lysed in 1x Laemmli buffer containing β-mercaptoethanol. The lysate was boiled at 95°C for 5 min. Proteins were separated by SDS–PAGE (Criterion TGX 4– 15%, Bio-Rad) and transferred to PVDF membranes using wet transfer. Blocking of membranes was carried out in 5% bovine serum albumin for 1 hr at room temperature, before incubation with primary antibodies diluted in 5% bovine serum albumin at 4°C for 16 h. Membranes were incubated with appropriate HRP-conjugated secondary antibodies (Cell Signaling Technologies, CST) for 1 hr at room temperature and developed using SuperSignal West Pico PLUS and SuperSignal West Femto substrates (ThermoFisher Scientific). The following primary antibodies were used: p65 (CST 8242), p(Ser^536^)-p65 (CST 3033), p(Thr^183^/Tyr^185^)-JNK (CST 4668), p(Ser^73^)-cJun (CST 9164), β-tubulin (Sigma-Aldrich T5201).

#### Viability assay

Cells were seeded at 8000 cells/well in a 96 well plate (Corning 353219) 2 days prior to experiment. At the time of experiment, a BreatheEasy seal was placed on top of the 96 well plate, and cells were placed into incubator with Optoplate-96 directly on top of BreatheEasy seal for 3 hours. Cells that were induced with light received one hour light OFF, followed by two hours light ON. Light inputs ranged from 0-6.84mW. After induction, cells were immediately trypsinized with 100 ul/well for 5 mins. Cells were then spun at 500g for 5 min. After spinning, supernatant was removed and cells were washed in PBS. After PBS wash, cells were resuspended in diluted Live/Dead stain (Thermo Scientific L10120) and stained according to manufacturer’s instructions. After washing, cells were run on BD LSRFortessa or BD LSRII.

### Computational Details

#### Analysis of RNA-seq data

Reads were aligned to the *Mus musculus* genome using the cloud service Bluebee designed for Lexogen Quantseq data. Briefly, the reads were trimmed using Bbduk and then aligned to the GRCm38 genome using STAR (*49*). After alignment, counts were generated using HTSeq-count (*50*). Raw counts data was then used with DESeq2 to generate log_2_FC and FDR p values (*51*). Genes with average raw counts ≤ 2 across all samples of interest were dropped from the analysis. log_2_FC RelA-CLASP induction was calculated for genes significantly regulated (FDR *p* < .05) by RelA-CLASP as such: log_2_FC (RelA-CLASP time t vs time 0) - log_2_FC (pm-LOVTRAP time t vs time 0), where t is either 1 or 2 hours of light induction. log_2_FC TNFɑ induction was defined as log_2_FC (pm-LOVTRAP time T vs time 0) for 1 or 2 hours of TNFɑ induction. log_2_FC induction is calculated from 5 replicates for each cell line and input.

#### Computational modeling

Modeling is as described in Chen 2020 for the two-state promoter model. Briefly, ordinary differential equations representing promoter kinetics were constructed with three state variables and six parameters. Parameters for k_on_, k_off_, and ɣ_1_ were sampled with uniform and London Hypercube sampling across of 0.002-2, 0.004-4, and .001-1, respectively. Parameters for β_0_ and β_1_ were set to 0.0032 and 4.92 for all simulations, respectively.

#### Image analysis

Microscopy images are analyzed for nuclear and cytoplasmic intensity using a custom Python script modified from nuclealyzer, which depends on StarDist, Scikit-image, and OpenCV (*52*– *54*). First, StarDist is used on nuclear BFP images to create masks of nuclei. Then, the cytoplasm is approximated by dilating the nuclear mask four times and subtracting a twice-dilated nuclear mask. Background of each image is estimated by expanding all nuclear masks in an image by 50 pixels, which approximates the cell radius, and then taking the mode of the intensity of the pixels which are not labeled by a mask. Finally, OpenCV is used to track centroids of the nuclear masks throughout the experiment. Nuclear/cytoplasmic enrichment (background subtracted) is calculated as (avg nuclear intensity - background intensity) / (avg cytoplasmic intensity - background intensity) for each cell. Images where blue light is on are dropped from analysis due to inability to isolate the nuclear marker in BFP images. Max Nuclear/cytoplasmic enrichment (background subtracted, norm to 0m) is calculated as follows: for each replicate and input, the frame with the maximum median nuclear/cytoplasmic enrichment is selected, and the nuclear/cytoplasmic enrichment for each cell present in that frame is normalized to the maximum median nuclear/cytoplasmic enrichment for the no light input in that replicate. Norm change in Nuclear/Cytoplasmic Enrichment is calculated as the median change in nuclear/cytoplasmic enrichment for each replicate divided by the maximum value of the median change in nuclear/cytoplasmic enrichment across RelA-mScarlet and RelA-CLASP cell lines.

#### Western Blot Quantification

Western blot band intensities (mean gray value) were quantified using ImageJ 1.53c. For determination of protein levels, blot background intensities are deducted from band intensities and background-corrected intensities are divided by β-tubulin levels for the same sample. Phospho-protein background-corrected, tubulin-normalized intensities are further divided by corresponding total protein levels, normalized to tubulin. RelA (65 kDa) level relative to NIH3T3 no light is calculated for each replicate as: (total RelA level (65 kDa)) /average(total RelA level (65 kDa)) where the average is taken across all NIH3T3 no light replicates. RelA (115 kDa) level relative to Rela-CLASP is calculated for each replicate as: (total RelA level (115 kDa)) /average(total RelA level (115 kDa)) where the average is taken across all Rela-CLASP no light replicates. p-Rela (65 kDa) level relative to TNF is calculated for each NIH3T3 replicate as: (p-RelA level (65 kDa))/(p-RelA level (65 kDa), NIH3T3 TNF). p-Rela (115 kDa) level relative to TNF is calculated for each RelA-CLASP replicate as: (p-RelA level (115 kDa))/(p-RelA level (115 kDa), RelA-CLASP TNF). p-JNK level relative to TNF is calculated for each replicate as: (p-JNK level (54 kDa)/(p-JNK level (54 kDa), TNF for same cell line). For S4B, cell line and input combinations were compared to the reference using a two-sample t-test with Holm-Sidak correction. For S4C-D, cell line and input combinations were compared to the reference using a one-sample t-test with Holm-Sidak correction.

#### Data processing

Data processing was done with custom-written Python, R, and Matlab scripts, which are available upon request.

## Acknowledgements

This work was supported by R01 GM119033-01A1 and a Chan-Zuckerberg Biohub gift (awarded to H.E-S.), R01AI127864 (awarded to A.H.), the National Defense Science & Engineering Graduate (NDSEG) Fellowship (awarded to L.C. O. and A.R.B.), Paul and Daisy Soros Fellowship for New Americans (awarded to L.C.O.), the National Science Foundation (NSF) Graduate Research Fellowship (GRF) (awarded to S.R.T.), and a postdoctoral fellowship from the Deutsche Forschungsgemeinschaft (DFG, German Research Foundation, 419234150) (awarded to S.L.). H.E-S. was an investigator in the Chan Zuckerberg Biohub and at Altos Labs at the time of research execution. This study was supported in part by the HDFCC Laboratory for Cell Analysis Shared Resource Facility through a grant from the NIH (P30CA082103). Data for this study were acquired at the Center for Advanced Light Microscopy at UCSF on the CSU-W1 Spinning Disk Confocal obtained with NIH S10 Shared Instrumentation grant (1S10OD017993-01A1). Sequencing was performed at the UCSF CAT, supported by UCSF PBBR, RRP IMIA, and NIH 1S10OD028511-01 grants. We thank Roy Wollman, Joao Fonseca, Helen Huang, and the El-Samad Lab for helpful discussions, Emily Wong and the Narlikar Lab for reagents, Zara Weinberg for assistance with cell tracking, image analysis, and RNA-seq, and Jesslyn Park for assistance with RNA-seq.

## Author Contributions

L.O.and H.E-S. conceived of the study and designed the methodology of the study. L.O. wrote code used to analyze the data. L.O. and S.L. analyzed the data. L.O., A.R.B., S.R.T, N.R., S.L., collected data and performed experiments. A.H. generated 3T3 *Nfkbia*^-/-^ *Nfkbib*^-/-^ *Nfkbie*^-/-^ *RelA*^-/-^ cells and provided the modified RelA construct. L.O., A.R.B., S.R.T., and N.R. constructed parts and cell lines. L.O. and H.E-S. prepared the original manuscript. All authors reviewed and edited the manuscript. L.O. and S.L. created the figures. L.O., A.H., and H.E-S. supervised the study. L.O. and H.E-S. coordinated the study. H.E-S. acquired the funding for the study.

## Data and Materials Availability

Further information and requests for resources and reagents should be directed to and will be fulfilled by the Lead Contact, Hana El-Samad. To request reagents, please submit a form to UCSF at https://ita.ucsf.edu/researchers/mta. All plasmids will also be deposited on Addgene and can be requested from there.

## References

1. R. Moreno, J.-M. Sobotzik, C. Schultz, M. L. Schmitz, Specification of the NF-κB transcriptional response by p65 phosphorylation and TNF-induced nuclear translocation of IKKε. Nucleic Acids Res. 38, 6029 (2010).

2. J.-P. Kruse, W. Gu, Modes of p53 Regulation. Cell. 137, 609–622 (2009).

3. H. Buss, A. Dörrie, M. L. Schmitz, E. Hoffmann, K. Resch, M. Kracht, Constitutive and Interleukin-1-inducible Phosphorylation of p65 NF-κB at Serine 536 Is Mediated by Multiple Protein Kinases Including IκB Kinase (IKK)-α, IKKβ, IKKϵ, TRAF Family Member-associated (TANK)-binding Kinase 1 (TBK1), and an Unknown Kinase and Couples p65 to TATA-binding Protein-associated Factor II31-mediated Interleukin-8 Transcription. J. Biol. Chem. 279, 55633–55643 (2004).

4. E. Batchelor, A. Loewer, C. Mock, G. Lahav, Stimulus-dependent dynamics of p53 in single cells. Mol. Syst. Biol. 7 (2011), doi:10.1038/msb.2011.20.

5. N. Yissachar, T. S. S. Fischler, A. A. Cohen, S. Reich-Zeliger, D. Russ, E. Shifrut, Z. Porat, N. Friedman, Dynamic response diversity of NFAT isoforms in individual living cells. Mol Cell. 49, 322–330 (2013).

6. A. Hoffmann, A. Levchenko, M. L. Scott, D. Baltimore, The IκB-NF-κB Signaling Module: Temporal Control and Selective Gene Activation. Science. 298, 1241–1245 (2002).

7. P. Hannanta-Anan, B. Y. Chow, Optogenetic Control of Calcium Oscillation Waveform Defines NFAT as an Integrator of Calcium Load. Cell Syst. 2, 283–288 (2016).

8. M. Son, A. G. Wang, H.-L. Tu, M. O. Metzig, P. Patel, K. Husain, J. Lin, A. Murugan, A. Hoffmann, S. Tay, NF-κB responds to absolute differences in cytokine concentrations. Sci. Signal. 14, 4382 (2021).

9. K. Lane, D. V. Valen, M. M. DeFelice, D. N. Macklin, T. Kudo, A. Jaimovich, A. Carr, T. Meyer, D. Pe’er, S. C. Boutet, M. W. Covert, Measuring Signaling and RNA-Seq in the Same Cell Links Gene Expression to Dynamic Patterns of NF-κB Activation. Cell Syst. 4, 458-469.e5 (2017).

10. A. Hafner, J. Stewart-Ornstein, J. E. Purvis, W. C. Forrester, M. L. Bulyk, G. Lahav, p53 pulses lead to distinct patterns of gene expression albeit similar DNA-binding dynamics. Nat. Struct. Mol. Biol. 2017 2410. 24, 840–847 (2017).

11. M. D. Harton, W. S. Koh, A. D. Bunker, A. Singh, E. Batchelor, p53 pulse modulation differentially regulates target gene promoters to regulate cell fate decisions. Mol. Syst. Biol. 15, e8685 (2019).

12. A. G. Goglia, M. Z. Wilson, S. G. Jena, J. Silbert, L. P. Basta, D. Devenport, J. E. Toettcher, A Live-Cell Screen for Altered Erk Dynamics Reveals Principles of Proliferative Control. Cell Syst. 10, 240-253.e6 (2020).

13. E. W. Martin, A. Pacholewska, H. Patel, H. Dashora, M. H. Sung, Integrative analysis suggests cell type–specific decoding of NF-κB dynamics. Sci. Signal. 13(2020), doi:10.1126/SCISIGNAL.AAX7195.

14. M. Z. Wilson, P. T. Ravindran, W. A. Lim, J. E. Toettcher, Tracing Information Flow from Erk to Target Gene Induction Reveals Mechanisms of Dynamic and Combinatorial Control. Mol. Cell. 67, 757-769.e5 (2017).

15. M. M. DeFelice, H. R. Clark, J. J. Hughey, I. Maayan, T. Kudo, M. V. Gutschow, M. W. Covert, S. Regot, NF-B signaling dynamics is controlled by a dose-sensing autoregulatory loop. Sci. Signal. 12, 3568 (2019).

16. S. Sen, Z. Cheng, K. M. Sheu, Y. H. Chen, A. Hoffmann, Gene regulatory strategies that decode the duration of NFκB dynamics contribute to LPS-versus TNF-specific gene expression. Cell Syst. 10, 169-182.e5 (2020).

17. S. Y. Chen, L. C. Osimiri, M. Chevalier, L. J. Bugaj, T. H. Nguyen, R. A. Greenstein, A. H. Ng, J. Stewart-Ornstein, L. T. Neves, H. El-Samad, Optogenetic Control Reveals Differential Promoter Interpretation of Transcription Factor Nuclear Translocation Dynamics. Cell Syst. 11, 336-353.e24 (2020).

18. H. Yumerefendi, D. J. Dickinson, H. Wang, S. P. Zimmerman, J. E. Bear, B. Goldstein, K. Hahn, B. Kuhlman, Control of protein activity and cell fate specification via light-mediated nuclear translocation. PLoS One. 10, e0128443 (2015).

19. H. Wang, M. Vilela, A. Winkler, M. Tarnawski, I. Schlichting, H. Yumerefendi, B. Kuhlman, R. Liu, G. Danuser, K. M. Hahn, LOVTRAP: An optogenetic system for photoinduced protein dissociation. Nat. Methods. 13, 755–758 (2016).

20. S. P. Heximer, H. Lim, J. L. Bernard, K. J. Blumer, Mechanisms governing subcellular localization and function of human RGS2. J Biol Chem. 276, 14195–14203 (2001).

21. J. P. Fonseca, A. R. Bonny, G. R. Kumar, A. H. Ng, J. Town, Q. C. Wu, E. Aslankoohi, S. Y. Chen, G. Dods, P. Harrigan, L. C. Osimiri, A. L. Kistler, H. El-Samad, A Toolkit for Rapid Modular Construction of Biological Circuits in Mammalian Cells. ACS Synth. Biol. 8, 2593–2606 (2019).

22. M. Bloom, S. Saksena, G. Swain, M. Behar, T. Yankeelov, A. Sorace, The effects of IKK-beta inhibition on early NF-kappa-B activation and transcription of downstream genes. Cell. Signal. 55, 17–25 (2019).

23. S. Kosugi, M. Hasebe, N. Matsumura, H. Takashima, E. Miyamoto-Sato, M. Tomita, H. Yanagawa, Six classes of nuclear localization signals specific to different binding grooves of importin alpha. J Biol Chem. 284, 478–485 (2009).

24. S. P. Mukherjee, M. Behar, H. A. Birnbaum, A. Hoffmann, P. E. Wright, G. Ghosh, Analysis of the RelA:CBP/p300 Interaction Reveals Its Involvement in NF-κB-Driven Transcription. PLOS Biol. 11, e1001647 (2013).

25. M. Urata, S. Kokabu, T. Matsubara, G. Sugiyama, C. Nakatomi, H. Takeuchi, S. Hirata-Tsuchiya, K. Aoki, Y. Tamura, Y. Moriyama, Y. Ayukawa, M. Matsuda, M. Zhang, K. Koyano, C. Kitamura, E. Jimi, A peptide that blocks the interaction of NF-κB p65 subunit with Smad4 enhances BMP2-induced osteogenesis. J. Cell. Physiol. 233, 7356–7366 (2018).

26. L. J. Bugaj, W. A. Lim, High-throughput multicolor optogenetics in microwell plates. Nat. Protoc. 2019 147. 14, 2205–2228 (2019).

27. L. J. Bugaj, A. J. Sabnis, A. Mitchell, J. E. Garbarino, J. E. Toettcher, T. G. Bivona, W. A. Lim, Cancer mutations and targeted drugs can disrupt dynamic signal encoding by the Ras-Erk pathway. Science. 361, 6405 (2018).

28. L. Verstrepen, T. Bekaert, T.-L. Chau, J. Tavernier, A. Chariot, R. Beyaert, TLR-4, IL-1R and TNF-R signaling to NF-κB: variations on a common theme. Cell. Mol. Life Sci. 2008 6519. 65, 2964–2978 (2008).

29. M. W. Covert, T. H. Leung, J. E. Gaston, D. Baltimore, Achieving Stability of Lipopolysaccharide-Induced NF-κB Activation. Science. 309, 1854–1857 (2005).

30. L. Ashall, C. A. Horton, D. E. Nelson, P. Paszek, C. V. Harper, K. Sillitoe, S. Ryan, D. G. Spiller, J. F. Unitt, D. S. Broomhead, D. B. Kell, D. A. Rand, V. Sée, M. R. H. White, Pulsatile stimulation determines timing and specificity of NF-kappa B-dependent transcription. Science. 324, 242 (2009).

31. S. Tay, J. J. Hughey, T. K. Lee, T. Lipniacki, S. R. Quake, M. W. Covert, Single-cell NF-κB dynamics reveal digital activation and analogue information processing. Nat. 2010 4667303. 466, 267–271 (2010).

32. Q. Zhang, S. Gupta, D. L. Schipper, G. J. Kowalczyk, A. E. Mancini, J. R. Faeder, R. E. C. Lee, NF-κB Dynamics Discriminate between TNF Doses in Single Cells. Cell Syst. 5, 638-645.e5 (2017).

33. C. O. Chavarría-Velázquez, A. C. Torres-Martínez, L. F. Montaño, E. P. Rendón-Huerta, TLR2 activation induced by H. pylori LPS promotes the differential expression of claudin-4,-6, -7 and -9 via either STAT3 and ERK1/2 in AGS cells. Immunobiology. 223, 38–48 (2018).

34. A. Müller, A. Hennig, S. Lorscheid, P. Grondona, K. Schulze-Osthoff, S. Hailfinger, D. Kramer, IκBζ is a key transcriptional regulator of IL-36–driven psoriasis-related gene expression in keratinocytes. Proc. Natl. Acad. Sci. 115, 10088–10093 (2018).

35. B. H. Rovin, J. A. Dickerson, L. C. Tan, C. A. Hebert, Activation of nuclear factor-κB correlates with MCP-1 expression by human mesangial cells. Kidney Int. 48, 1263–1271 (1995).

36. F. Luan, X. Li, X. Cheng, L. Huangfu, J. Han, T. Guo, H. Du, X. Wen, J. Ji, TNFRSF11B activates Wnt/β-catenin signaling and promotes gastric cancer progression. Int. J. Biol. Sci. 16, 1956 (2020).

37. M. Neumann, M. Naumann, Beyond IκBs: alternative regulation of NF-KB activity. FASEB J. 21, 2642–2654 (2007).

38. K. A. Ngo, K. Kishimoto, J. Davis-Turak, A. Pimplaskar, Z. Cheng, R. Spreafico, E. Y. Chen, A. Tam, G. Ghosh, S. Mitchell, A. Hoffmann, Dissecting the Regulatory Strategies of NF-κB RelA Target Genes in the Inflammatory Response Reveals Differential Transactivation Logics. Cell Rep. 30, 2758-2775.e6 (2020).

39. P. Rao, M. S. Hayden, M. Long, M. L. Scott, A. P. West, D. Zhang, A. Oeckinghaus, C. Lynch, A. Hoffmann, D. Baltimore, S. Ghosh, IκBβ acts to inhibit and activate gene expression during the inflammatory response. Nat. 2010 4667310. 466, 1115–1119 (2010).

40. C. Genolini, B. Falissard, KmL: k-means for longitudinal data. Comput. Stat. 2009 252. 25, 317–328 (2009).

41. A. P. Gasch, P. T. Spellman, C. M. Kao, O. Carmel-Harel, M. B. Eisen, G. Storz, D. Botstein, P. O. Brown, Genomic expression programs in the response of yeast cells to environmental changes. Mol Biol Cell. 11, 4241–4257 (2000).

42. N. Vardi, S. Levy, Y. Gurvich, T. Polacheck, M. Carmi, D. Jaitin, I. Amit, N. Barkai, Sequential feedback induction stabilizes the phosphate starvation response in budding yeast. Cell Rep. 9, 1122–1134 (2014).

43. A. Adelaja, B. Taylor, K. M. Sheu, Y. Liu, S. Luecke, A. H. Correspondence, Six distinct NFκB signaling codons convey discrete information to distinguish stimuli and enable appropriate macrophage responses. Immunity. 54, 916-930.e7 (2021).

44. M. V. Gutschow, J. C. Mason, K. M. Lane, I. Maayan, J. J. Hughey, B. T. Bajar, D. N. Amatya, S. D. Valle, M. W. Covert, Combinatorial processing of bacterial and host-derived innate immune stimuli at the single-cell level. httpsdoiorg101091mbcE18-07-0423. 30, 282–292 (2019).

45. S. Zambrano, A. Loffreda, E. Carelli, G. Stefanelli, F. Colombo, E. Bertrand, C. Tacchetti, A. Agresti, M. E. Bianchi, N. Molina, D. Mazza, First Responders Shape a Prompt and Sharp NF-κB-Mediated Transcriptional Response to TNF-α. iScience. 23, 101529 (2020).

46. A. E. Davies, M. Pargett, S. Siebert, T. E. Gillies, Y. Choi, S. J. Tobin, A. R. Ram, V. Murthy, C. Juliano, G. Quon, M. J. Bissell, J. G. Albeck, Systems-Level Properties of EGFR-RAS-ERK Signaling Amplify Local Signals to Generate Dynamic Gene Expression Heterogeneity. Cell Syst. 11, 161-175.e5 (2020).

47. M. E. Lee, W. C. DeLoache, B. Cervantes, J. E. Dueber, A highly characterized yeast toolkit for modular, multipart assembly. ACS Synth Biol. 4, 975–986 (2015).

48. O. S. Thomas, M. Hörner, W. Weber, A graphical user interface to design high-throughput optogenetic experiments with the optoPlate-96. Nat. Protoc. 2020 159. 15, 2785–2787 (2020).

49. A. Dobin, C. A. Davis, F. Schlesinger, J. Drenkow, C. Zaleski, S. Jha, P. Batut, M. Chaisson, T. R. Gingeras, STAR: ultrafast universal RNA-seq aligner. Bioinformatics. 29, 15 (2013).

50. S. Anders, P. T. Pyl, W. Huber, HTSeq—a Python framework to work with high-throughput sequencing data. Bioinformatics. 31, 166 (2015).

51. M. I. Love, W. Huber, S. Anders, Moderated estimation of fold change and dispersion for RNA-seq data with DESeq2. Genome Biol. 2014 1512. 15, 1–21 (2014).

52. Z. Weinberg, weinberz/nuclealyzer: Public Release (Zenodo, 2022; https://zenodo.org/record/6366817).

53. M. Weigert, U. Schmidt, R. Haase, K. Sugawara, G. Myers, Star-convex Polyhedra for 3D Object Detection and Segmentation in Microscopy. IEEE Winter Conf. Appl. Comput. Vis. WACV (2020) (available at https://github.com/).

54. U. Schmidt, M. Weigert, C. Broaddus, G. Myers, Cell Detection with Star-convex Polygons. Lect. Notes Comput. Sci. Subser. Lect. Notes Artif. Intell. Lect. Notes Bioinforma. 11071 LNCS, 265–273 (2018).

