## Supplemental Figures and Captions for "Optogenetic control of RelA reveals effect of transcription factor dynamics on downstream gene expression"

### Figure S1: Optimization and modularity of mammalian CLASP

A) Schematic of three-plasmid CLASP system. Parental cell lines can be transduced with two plasmids at once via lentivirus, one expressing pm-LOVTRAP and one expressing a nuclear marker, thereby creating a chassis cell line. The chassis cell line can then be transduced with a variety of Zdk1-cargo-yeLANS plasmids, where "cargo" can be any transcription factor. B) Images of chassis cell lines. Chassis cell lines created in HEK293T, MCF10A, and 3T3 *Nfkbia*$^{-/-}$ *Nfkbib*$^{-/-}$ *Nfkbie*$^{-/-}$ *RelA*$^{-/-}$ are shown. IRFP (false colored yellow) and BFP (false colored blue) channels are merged. IRFP shows pm-LOVTRAP localization and BFP is used as a nuclear marker. Scale bar is 10 μm in all images. C) Images of NFAT1-CLASP response to blue light. (Top panels) RFP image (false colored red) showing Zdk1-NFAT1-mScarlet-yeLANS. (Bottom panels) IRFP (false colored yellow) and BFP (false colored blue) channels are merged. IRFP shows pm-LOVTRAP localization and BFP is used as a nuclear marker. Cells are induced with 15 min of blue light between images. Scale bar is 10 μm. D) Images of p53 K305N-CLASP response to blue light. (Top panels) RFP image (false colored red) showing Zdk1-p53 K305N-mScarlet-yeLANS. (Bottom panels) IRFP (false colored yellow) and BFP (false colored blue) channels are merged. IRFP shows pm-LOVTRAP localization and BFP is used as a nuclear marker. Cells are induced with 10 min blue light between images. Scale bar is 10 μm.

### Figure S2: RelA-CLASP responds to light duration and intensity

A) Nuclear/cytoplasmic enrichment of RelA-CLASP as a function of time in response to different durations of blue light. Light input turns on at 0 minutes as shown on x-axis. Data are from 3 replicates with 553-665 cells tracked per input. Nuclear/Cytoplasmic Enrichment for each input is normalized to the value measured in the frame preceding light on. B) Nuclear/cytoplasmic enrichment of RelA-CLASP as a function of time in response to different intensities of blue light. Light input is shown above plot; light turns on at 5 minutes. Data are from 3 replicates with 152-209 cells tracked per input. Asterisk denotes the 9-minute data point for 1.26 mW light, which has data only from 1 replicate. Nuclear/Cytoplasmic Enrichment for each input is normalized to the value measured at 0 minutes.

### Figure S3: RelA-CLASP induces gene expression in response to blue light

A) Gene Ontology (GO) term plot for genes significantly (FDR $p < .05$, n=1209) regulated by 1ng/mL TNFα in pm-LOVTRAP cells. Benjamini-Hochberg adjusted p value is plotted on the y-axis and the number of overlapping genes for each Gene Ontology term is shown through color. A subset of GO terms is shown. B) GO term plot for genes significantly (FDR $p < .05$) regulated by 1 ng/mL TNFα in pm-LOVTRAP cells which are not significantly (FDR $p > .05$) regulated by RelA-CLASP following a constant light input (n= 559). Data is plotted as described in A. A subset of GO terms is shown. C) Dose response of viability in response to blue light intensities for 3T3 *Nfkbia*$^{-/-}$ *Nfkbib*$^{-/-}$ *Nfkbie*$^{-/-}$ *RelA*$^{-/-}$, pm-LOVTRAP, and RelA-CLASP cell lines. Light input is delivered as shown in figure 3A. Viability is plotted as the fraction of cells that are alive. Black triangle denotes median viability across 9 replicates; gray circles show viability for each replicate. D) Line plot of Figure 3E heatmap. $\log_2$FC RelA-CLASP induction, normalized to constant max is calculated by dividing $log_2$FC RelA-CLASP induction for a given gene by the maximum $log_2$FC RelA-CLASP induction for that gene. Longitudinal gene trajectories are grouped into 6 clusters using k-means longitudinal clustering.

### Figure S4: RelA-CLASP and JNK are not phosphorylated in response to blue light

A) Western blots of total RelA, total JNK, phospho-RelA (S536), and phospho-JNK protein levels in RelA-CLASP, pm-LOVTRAP, and NIH3T3 cells in response to light and TNFα induction. Cells are induced with either 30 min 1.38 mW blue light or 30 min 1 ng/ml TNFα. β-tubulin is shown as a loading control below the respective blots. (-) denotes no input, L denotes light input, and T denotes TNFα input. B) Quantification of total RelA Western Blots at 65 kDa and 115 kDa for RelA-CLASP, pm-LOVTRAP, and NIH3T3 cell lines. (Left panel) Relative RelA protein levels at endogenous weight (65 kDa). (Right panel) Relative RelA protein levels at Zdk1-RelA-mScarlet-yeLANS weight (115 kDa). C) Quantification of phospho-RelA (S536) Western Blots at 65 kDa and 115 kDa. (Left panel) Relative phospho-RelA (S536) protein levels at endogenous weight (65 kDa) for NIH3T3 cell line in response to no light and TNFα induction. (Right panel) Relative phospho-RelA (S536) protein levels at Zdk1-RelA-mScarlet-yeLANS weight (115 kDa) for RelA-CLASP cell line in response to no light, light, and TNFα induction. D) Quantification of phospho-JNK Western Blots at 54 kDa. (Left panel) Relative p-JNK protein levels for NIH3T3 cell line in response to no light and TNFα induction. (Right panel) Relative p-JNK protein levels for RelA-CLASP cell line in response to no light, light, and TNFα induction. For panels B-D, barplots represent mean value for each cell line and input, error bars represent standard deviation, and detailed description of normalization is provided in Methods. ns denotes adjusted $p > .05$, * denotes adjusted $p < .05$, ** denotes adjusted $p < .01$, *** denotes adjusted $p < .001$; all *p*-values from t-test with Holm-Sidak correction.

### Figure S5: Parameter regimes that determine response to constant and pulsed inputs for early genes using simple promoter model

A) Scatter plots of $k_{on}$ and $k_{off}$ parameter values simulated with the simple promoter model. (Left panel) All parameter sets simulated with a constant RelA-CLASP input are plotted in black. (Right panel) Parameter sets that generate an early gene response are plotted in dark gray. B) Scatter plots of $k_{on}$ and $\gamma_1$ parameter values simulated with the simple promoter model. Points are plotted and colored as described in panel A. C) Scatter plots of $k_{off}$ and $\gamma_1$ parameter values simulated with the simple promoter model. Points are plotted and colored as described in panel A. D) Normalized simulated $log_2$FC RelA-CLASP induction as a function of time for a subset (n=100) of early gene parameter sets in response to constant RelA-CLASP input. Simulated $log_2$FC RelA-CLASP induction, normalized to constant max is calculated as the simulated $log_2$FC RelA-CLASP induction values at all timepoints divided by the maximum simulated $log_2$FC RelA-CLASP induction measured at 0, 120, or 180 min of two-hour constant light simulation for each parameter set. Dark colored bands show 25th-75th percentile of normalized simulated $log_2$FC RelA-CLASP induction; light colored bands show 0-100th percentile of the same quantity. Dashed lines show sampling times for 0 hr, 1 hr, and 2 hr cumulative light induction time, which correspond to RNA-seq sample collection. E) Clustering of response to pulsed input for parameter sets which recapitulate the early gene response to a constant light input. (All panels) Plots of normalized simulated $log_2FC$ RelA-CLASP induction as a function of cumulative light induction time, clustered with longitudinal k-means clustering. (Left panels) k=3 clusters. Colors shown above plots correspond to clusters shown in Figure 4D. (Right panels) k=5 clusters. F) Scatter plots of parameter values for $k_{on}$ and $k_{off}$ that yield different dynamics in response to a pulsed RelA-CLASP input. All parameter sets shown generate an early response to a constant input. Colors map to clusters shown in Figure 4D. G) Scatter plots of parameter values for $k_{on}$ and $\gamma_1$ that yield varied dynamics in response to a pulsed RelA-CLASP input. All parameter sets shown generate an early response to a constant input in log space. Colors map to clusters shown in Figure 4D. H) Normalized simulated $log_2FC$ RelA-CLASP induction vs time in response to pulsed TF input for a subset of parameter sets in 4D. Simulated $log_2FC$ RelA-CLASP induction, normalized to constant max is calculated as described in D. Color of bands corresponds to clustering as shown in Figure 4D. Simulated $log_2FC$ RelA-CLASP induction is plotted for 100 parameter sets for each cluster. Dark colored bands show 25th-75th percentile of normalized simulated $log_2FC$ RelA-CLASP induction values; light colored bands show 0-100th percentile of the same quantity. Dashed lines show sampling times for 0 hr, 1 hr, and 2 hr cumulative light induction time.

### Figure S6: Parameter regimes that determine response to constant and pulsed inputs for proportional genes using simple promoter model.

A) Scatter plots of $k_{on}$ and $k_{off}$ parameter values simulated with the simple promoter model. (Left panel) All parameter sets simulated with a constant RelA-CLASP input are plotted in black. (Right panel) Parameter sets that generate a proportional gene response are plotted in light gray. B) Scatter plots of $k_{on}$ and $\gamma_1$ parameter values simulated with the simple promoter model. Points are plotted and colored as described in panel A. C) Scatter plots of $k_{off}$ and $\gamma_1$ parameter values simulated with the simple promoter model. Points are plotted and colored as described in panel A. D) Normalized simulated $log_2FC$ RelA-CLASP induction as a function of time for a subset (n=100) of proportional gene parameter sets in response to constant RelA-CLASP input. Simulated $log_2FC$ RelA-CLASP induction, normalized to constant max is calculated as the simulated $log_2FC$ RelA-CLASP induction values at all timepoints divided by the maximum simulated $log_2FC$ RelA-CLASP induction measured at 0, 120, or 180 min of constant input simulation for each parameter set. Dark colored bands show 25th-75th percentile of normalized simulated $log_2FC$ RelA-CLASP induction; light colored bands show 0-100th percentile of the same quantity. Dashed lines show sampling times for 0h, 1h, and 2h cumulative light induction time, which correspond to RNA-seq sample collection. E) Clustering of response to pulsed input for parameter sets which recapitulate the proportional gene response to a two-hour constant light input. (All panels) Plots of normalized simulated $log_2FC$ RelA-CLASP induction as a function of cumulative light induction time, clustered with longitudinal k-means clustering. (Left panels) k=2 clusters. Colors shown above plots correspond to clusters shown in Figure 5C. (Right panels) k=6 clusters. F) Scatter plots of parameter values for $k_{on}$ and $k_{off}$ that yield varied dynamics in response to a pulsed RelA-CLASP input. All parameter sets shown generate a proportional response to a constant input. Colors map to clusters shown in Figure 5C. G)

Scatter plots of parameter values for $k_{on}$ and $\gamma_1$ that yield varied dynamics in response to a pulsed RelA-CLASP input. All parameter sets shown generate a proportional response to a constant input. Colors map to clusters shown in Figure 5C. H) Normalized simulated $log_2FC$ RelA-CLASP induction vs time in response to pulsed TF input for a subset of parameter sets in 5C. Simulated $log_2FC$ RelA-CLASP induction, normalized to constant max is calculated as described in D. Color corresponds to clustering shown in Figure 5C. Simulated $log_2FC$ RelA-CLASP induction is plotted for 100 parameter sets for each cluster. Dark colored bands show 25th-75th percentile of normalized simulated $log_2FC$ RelA-CLASP induction; light colored bands show 0-100th percentile of the same quantity. Dashed lines show sampling times for 0h, 1h, and 2h cumulative light induction time.

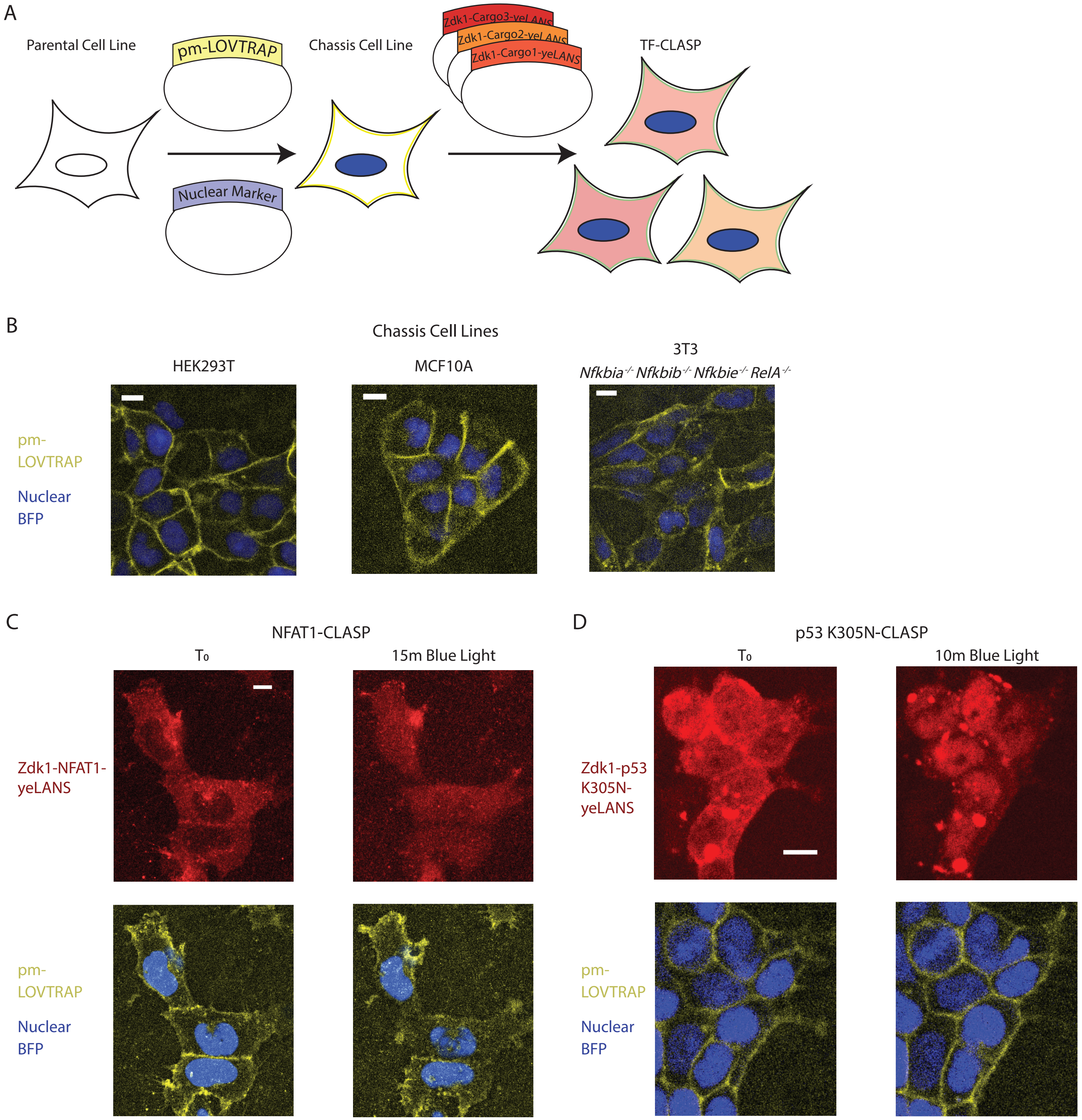
A
Parental Cell Line
pm-LOVTRAP
Nuclear Marker
Chassis Cell Line
Zdk1-Cargo3-yeLANS
Zdk1-Cargo2-yeLANS
Zdk1-Cargo1-yeLANS
TF-CLASP
B
Chassis Cell Lines
HEK293T
MCF10A
3T3
Nfkbia-/- Nfkbib-/- Nfkbie-/- RelA-/-
pm-LOVTRAP
Nuclear BFP
C
NFAT1-CLASP
T0
15m Blue Light
Zdk1-NFAT1-yeLANS
pm-LOVTRAP
Nuclear BFP
D
p53 K305N-CLASP
T0
10m Blue Light
Zdk1-p53 K305N-yeLANS
pm-LOVTRAP
Nuclear BFP

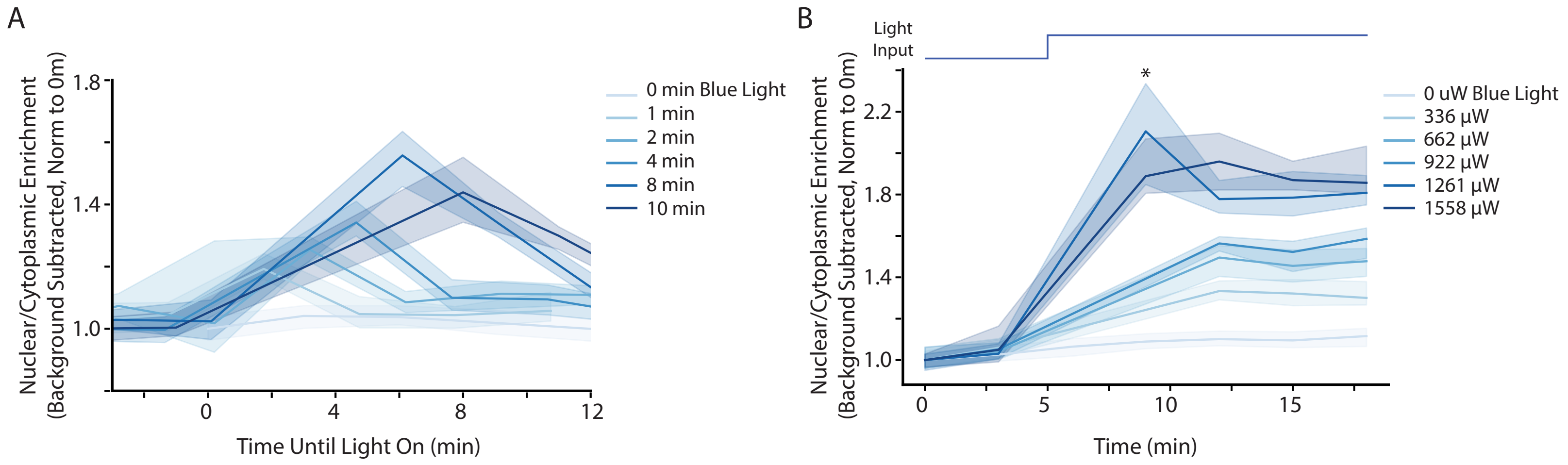

A
Nuclear/Cytoplasmic Enrichment (Background Subtracted, Norm to 0m)
1.8
1.4
1.0
0
4
8
12
Time Until Light On (min)
0 min Blue Light
1 min
2 min
4 min
8 min
10 min
B
Light Input
*
Nuclear/Cytoplasmic Enrichment (Background Subtracted, Norm to 0m)
2.2
1.8
1.4
1.0
0
5
10
15
Time (min)
0 uW Blue Light
336 μW
662 μW
922 μW
1261 μW
1558 μW

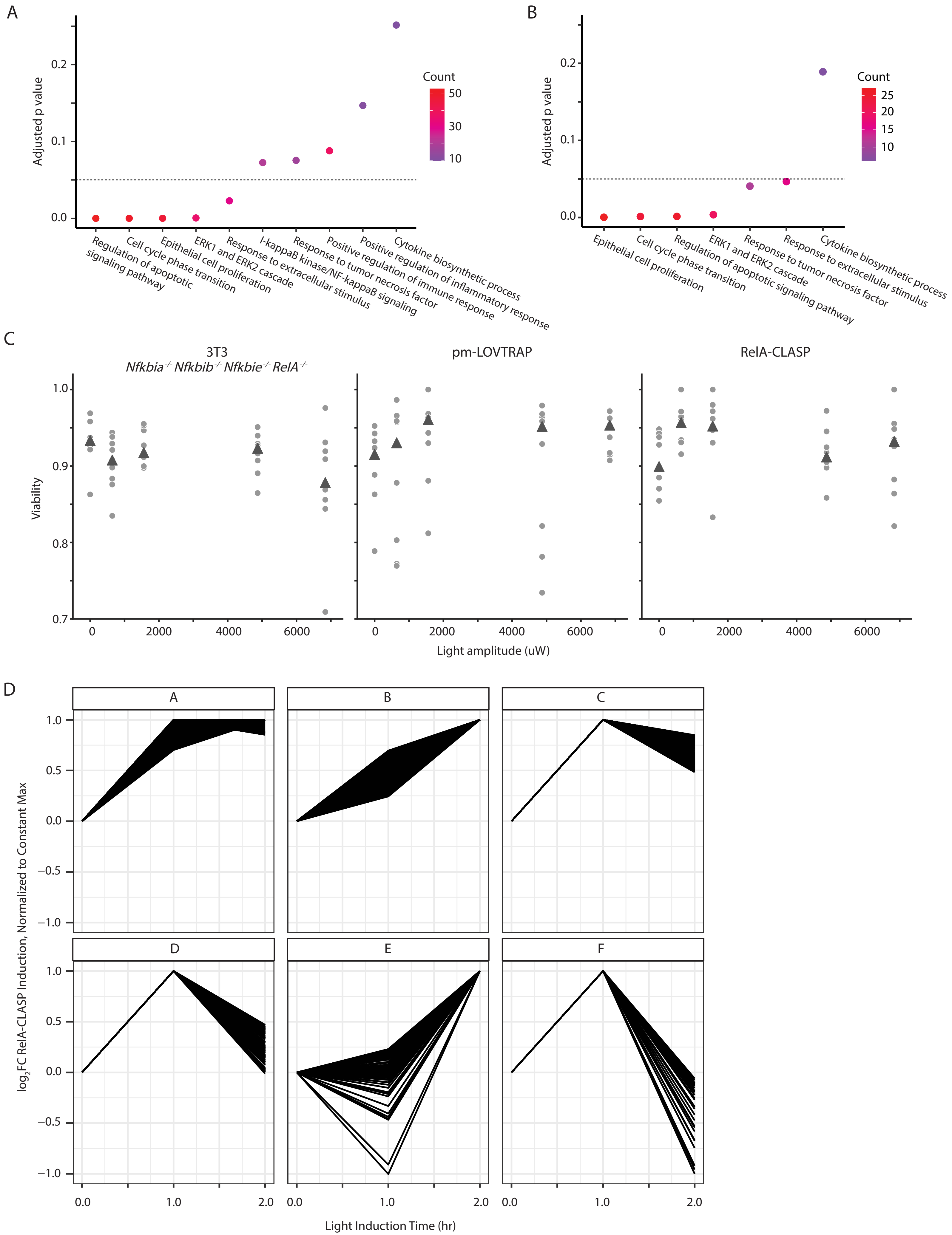
A
Adjusted p value
0.0
0.1
0.2
Count
50
30
10
Regulation of apoptotic signaling pathway
Cell cycle phase transition
Epithelial cell proliferation
ERK1 and ERK2 cascade
Response to extracellular stimulus
I-kappaB kinase/NF-kappaB signaling
Response to tumor necrosis factor
Positive regulation of immune response
Positive regulation of inflammatory response
Cytokine biosynthetic process
B
Adjusted p value
0.0
0.1
0.2
Count
25
20
15
10
Epithelial cell proliferation
Cell cycle phase transition
Regulation of apoptotic signaling pathway
ERK1 and ERK2 cascade
Response to tumor necrosis factor
Response to extracellular stimulus
Cytokine biosynthetic process
C
3T3
Nfkbia-/- Nfkbib-/- Nfkbie-/- RelA-/-
pm-LOVTRAP
RelA-CLASP
Viability
0.7
0.8
0.9
1.0
0
2000
4000
6000
Light amplitude (uW)
D
A
B
C
D
E
F
log2FC RelA-CLASP Induction, Normalized to Constant Max
−1.0
−0.5
0.0
0.5
1.0
0.0
1.0
2.0
Light Induction Time (hr)

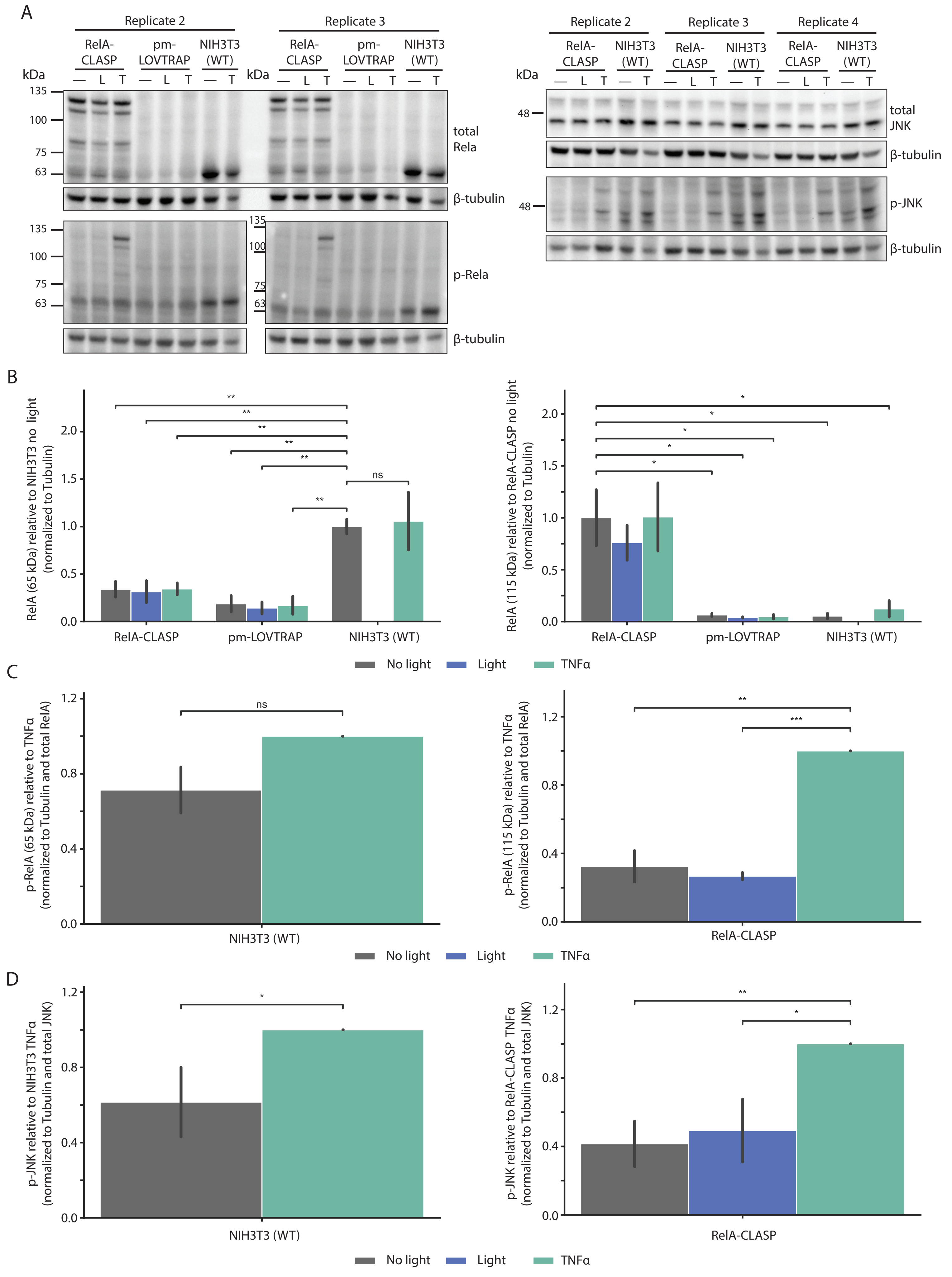
A
Replicate 2
Replicate 3
RelA-CLASP
pm-LOVTRAP
NIH3T3 (WT)
kDa
— L T
135
100
75
63
total Rela
β-tubulin
p-Rela
Replicate 4
48
total JNK
p-JNK
B
RelA (65 kDa) relative to NIH3T3 no light (normalized to Tubulin)
RelA (115 kDa) relative to RelA-CLASP no light (normalized to Tubulin)
**
*
ns
C
p-RelA (65 kDa) relative to TNFα (normalized to Tubulin and total RelA)
p-RelA (115 kDa) relative to TNFα (normalized to Tubulin and total RelA)
***
D
p-JNK relative to NIH3T3 TNFα (normalized to Tubulin and total JNK)
p-JNK relative to RelA-CLASP TNFα (normalized to Tubulin and total JNK)
No light
Light
TNFα

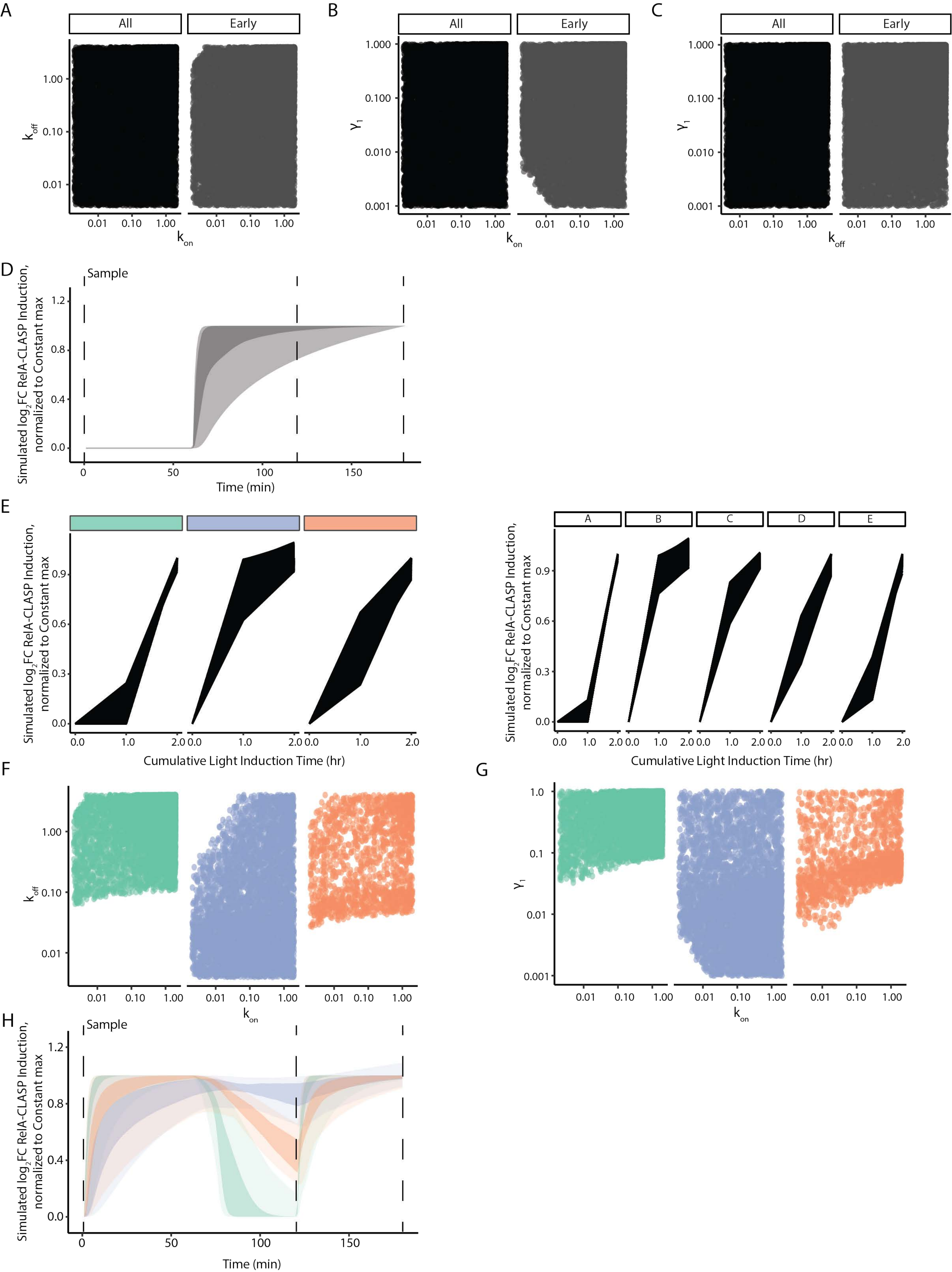
A
All
Early
1.00
0.10
0.01
$k_{off}$
0.01
0.10
1.00
$k_{on}$
B
$\gamma_1$
1.000
0.100
0.010
0.001
C
D
Sample
Simulated $log_2$FC RelA-CLASP Induction, normalized to Constant max
1.2
0.8
0.4
0.0
0
50
100
150
Time (min)
E
0.9
0.6
0.3
Cumulative Light Induction Time (hr)
0.0
1.0
2.0
A
B
C
D
E
F
G
1.0
0.1
0.01
H

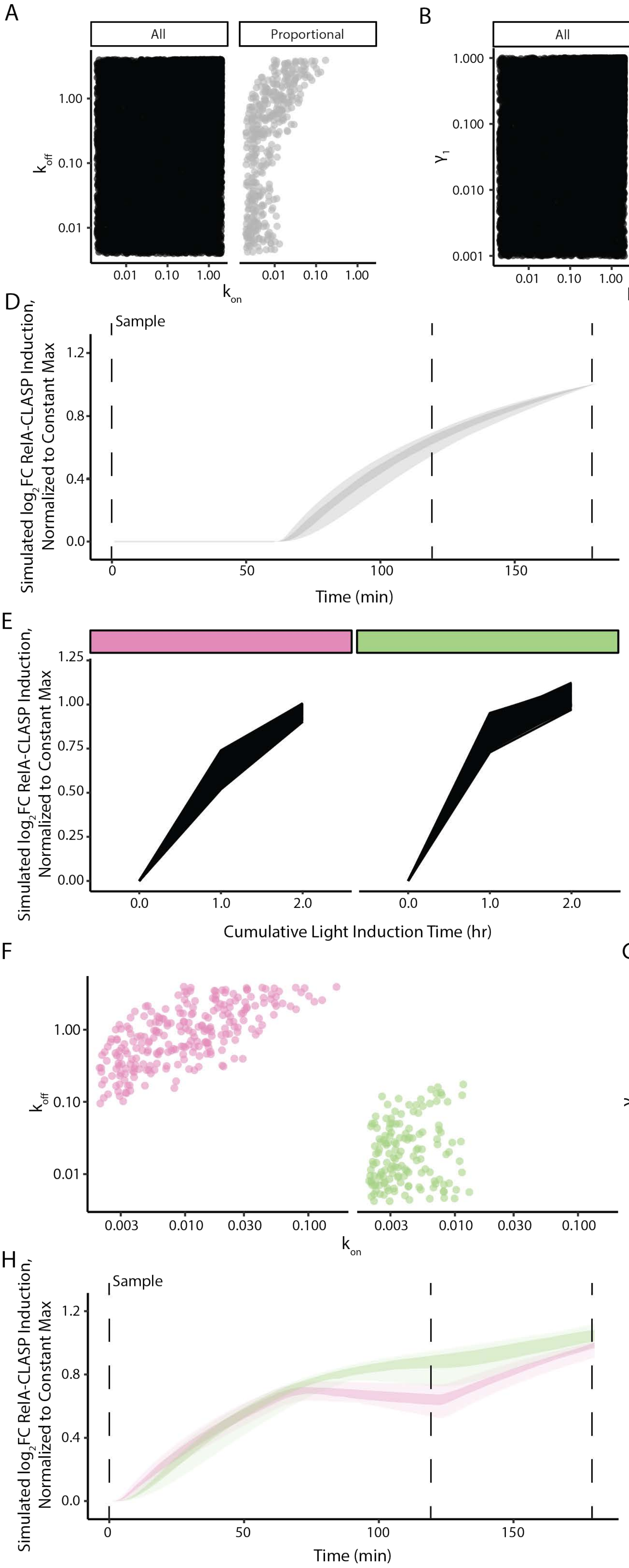
A
All
Proportional
$k_{off}$
$k_{on}$
B
$\gamma_1$
C
D
Sample
Simulated $log_2$FC RelA-CLASP Induction, Normalized to Constant Max
Time (min)
E
A
B
C
D
E
F
Cumulative Light Induction Time (hr)
F
G
H
